# 4polar3D : Single molecule 3D orientation imaging of dense actin networks using ratiometric polarization splitting

**DOI:** 10.1101/2025.07.13.664601

**Authors:** Charitra S. Senthil Kumar, Cesar A. Valades Cruz, Miguel Sison, Arturo G. Vesga, Javier Rey-Barroso, Valentina Curcio, Luis A. Alemán-Castañeda, Miguel A. Alonso, Renaud Poincloux, Manos Mavrakis, Sophie Brasselet

## Abstract

Single Molecule Orientation and Localization Microscopy (SMOLM) aims at simultaneously measuring the position and orientation of single molecules, generating orientation-encoded super-resolved images by estimating both their 3D mean orientation and the extent of their angular fluctuations (wobble). Most existing SMOLM approaches rely on the engineering of single molecules’ point spread functions, which requires complex optical setups and long computational times that can be an obstacle in dense cellular environments with high detection density and challenging imaging conditions. In this work, we propose a simpler and effective method named 4polar3D, based on the estimation of single molecule intensities projected onto four polarized channels with controlled numerical apertures. This strategy enables 3D orientation measurements of single molecules in addition to their 2D localization in a fast processing step. We demonstrate that 4polar3D can resolve nanoscale molecular organization in whole cells’ crowded structures, uncovering 3D-oriented actin filament networks in densely packed lamellipodia and podosomes.

## Introduction

Single Molecule Orientation and Localization Microscopy (SMOLM) emerged a decade ago with the aim of acheiving localization and orientation imaging of isolated fluorescent molecular dipoles^1,2^. Resolving single molecules’ (SMs) orientation in addition to their spatial position enables the investigation of the nanoscale 3D organization of proteins, which is crucial to understanding a wide range of biological processes from immunology to mechabiology. Molecular orientation plays indeed a critical role in biological function, not only at the level of proteins’ conformational changes, but also in the context of their collective structural organization. For example the orientation of actin filaments in the cell cytoskeleton, whose structure is typically inaccessible to standard single molecule localization microscopy (SMLM) due to their high molecular density, can be better probed using orientation-resolved techniques. Since the point spread function (PSF) of single molecules imaged with a high numerical aperture (NA) microscope encodes information on their orientation, most SMOLM strategies rely on analyzing the shape of engineered PSFs, typically modified using phase or polarization masks placed at the pupil plane of the microscope objective^3–9^. PSF engineering enables precise estimation of both the average orientation of SMs as well as their rotational dynamics referred to as wobble, which reflects the extent of their angular fluctuations over tens of milliseconds. This approach requires however advanced optimization and retrieval strategies^2^, such as PSF basis decomposition^7,10^, maximum likelihood estimation^11–13^ or deep learning approaches^14,15^. Despite the potential of PSF engineering for SMOLM, it has been predominantly applied to in vitro systems^11,16,17^ rather than to complex, densely packed cellular environments, which typically requires a very high number of localizations over large field of views. PSF engineering approaches have several inherent limitations, namely the complexity of microscope setups to be implemented, the use of enlarged PSFs that decrease the accessible detection density, the need to correct for phase and polarization aberrations^18–21^, and the substantial computational complexity associated with parameters’ retrieval, often resulting in slow data analysis procedures.

An alternative strategy to PSF engineering relies on the ratiometric intensity estimation of the signal from SMs polarized onto different polarization channels, in addition to standard 2D localization. By projecting the fluorescence signal onto two or four polarization channels, it is possible to estimate SMs’ anisotropy in isotropic environments^22^ or to determine their 2D orientation projected in the sample plane^23–26^. Extending this approach to 3D orientation estimation is not straightforward, as polarization is generally manipulated in the transverse optical plane to the propagation direction. So far, 3D orientation determination using ratiometric polarization-based methods has been achieved only indirectly, supposing that the molecules are fixed or have a known wobble^27,28^, involving sequential measurements with multiple directions of illumination^29^, or fitting polarized PSFs^13,26,30^ at the price of an increase in computation complexity. Among the least computationally demanding strategies, ratiometric polarization-based estimation seems highly appropriate for its relatively low sensitivity to microscope’s distorsions, its decrease in processing time of SMOLM, and its capacity to access information from large data sets in crowded environments.

In this work, we propose a simple, yet powerful scheme for measuring 3D orientation and wobble of SMs using a four-polarization ratiometric approach that relies solely on intensity estimations across distinct polarization channels. This scheme combines polarization splitting to extract in-plane orientation information with selective detection in the pupil plane of the microscope, to access off-plane orientation information. The approach capitalizes on the fact that the angular emission pattern radiated by a molecular dipole is stronlgy dependent on its off-plane tilt angle^31^. Specifically, when the dipole is tilted off-plane (i.e., perpendicular to the sample plane), the border region of the pupil plane, corresponding to the most tilted radiation wave vectors, is brighter than the central region (Fig. 1a, left). Inversely, when the dipole lies in the sample plane, the central region is brighter (Fig. 1a, right).

**Fig. 1.**
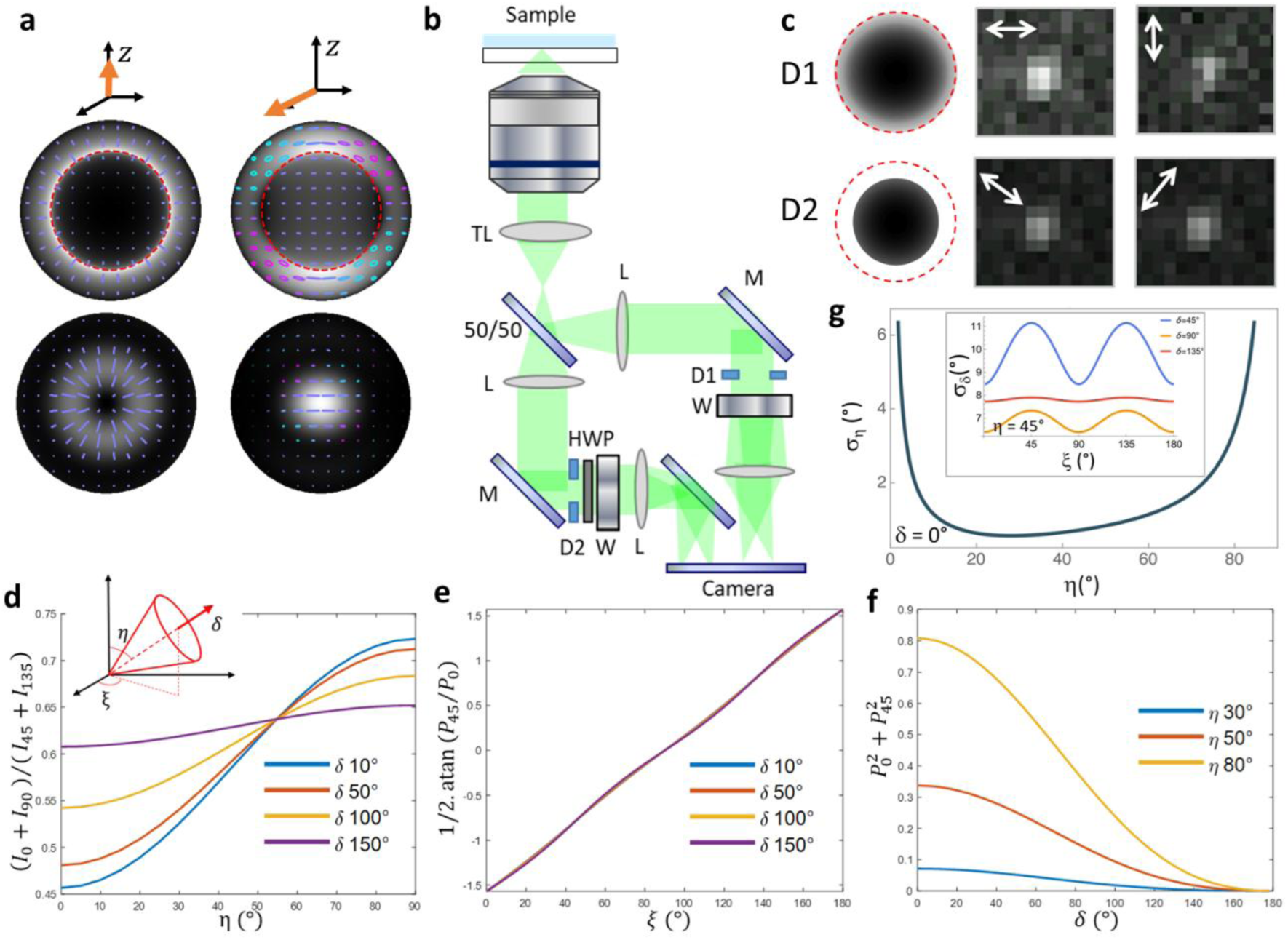
Principle of 4polar3D. **a** Intensity and polarization distributions in the pupil plane (top) and image plane (bottom) of a dipole (orange arrow) oriented along the longitudinal axis z (left) and the in-plane axis x (right). The intensity pattern (black and white image) is used as a background and the local polarization is drawn with a shape and color code representing its ellipticity (cyan: left handedness, magenta: right handedness). The dashed red circle corresponds to the super critical angle emission. **b** Schematic representation of the 4polar3D setup. TL: tube lens, 50/50: non-polarizing beam splitter, L: lenses, M: mirrors, D1 and D2: diaphragms, W: Wollaston prism, HWP: half wave plate. **c** Example of experimental PSFs from a single molecule along the four polarized channels (represented by the white arrow) (pixel size 130 nm). The (0°,90°) and (45°,135°) channels are measured respectively at NA = 1.1 (diaphragm D1) and 1.3 (diaphragm D2). **d** Dependence of the intensity ratio (𝐼_0_ + 𝐼_90_)/(𝐼_45_ + 𝐼_135_) on 𝜂 for different values of δ. Inset : definition of the angles ξ, 𝜂, 𝛿, the red arrow defines the mean direction of the single molecule wobbling cone. **e** Dependence of 1/2. atan 𝑃_45_/𝑃_0_ on ξ for different values of δ. **f** Dependence of 𝑃_0_^2^ + 𝑃_45_^2^ on δ for different values of η. **g** Cramér Rao lower bound on the expectation of the 𝜂 parameter, for ξ = 0° and a fixed dipole (𝛿 = 0°). Conditions used : total intensity 5 000 photons, 𝑛_0_ = 1.33, 𝑛_1_ = 1.515, 𝑧_0_ = 𝑧_1_ = 0, 𝑁𝐴_𝑙𝑜𝑤_ = 1.04, 𝑁𝐴_ℎ𝑖𝑔ℎ_ = 1.3 (under critical angle condition, 𝑁𝐴 ratio 0.8). Inset : CRLB on 𝛿 calculated as a function of ξ, for different values of 𝛿, for 𝜂 = 45°.

To acheive 3D orientation estimation, the method thus combines four polarization splitting channels with different effective numerical apertures (𝑁𝐴s). First, the signal is split onto two channels of different 𝑁𝐴s, respectively low 𝑁𝐴 (𝑁𝐴_𝑙𝑜𝑤_) and high 𝑁𝐴 (𝑁𝐴_ℎ𝑖𝑔ℎ_) using two different diaphragms, D1 and D2 respectively (Fig. 1b,c). Each of these two unpolarized channels is then split into two linear polarizations, 0°/90° and 45°/135° respectively, with respect to the horizontal axis of the sample plane, as introduced previously,^25^ in a relatively compact and simple implementation (Fig. 1b). This method, named 4polar3D, combines the benefits of transverse 4-polarization splitting and pupil amplitude splitting (e.g. NA filtering) to determine the azimuth angle (i.e., in-plane orientation), polar angle (i.e. off-plane orientation) to within one sign, and the wobble angle of SMs, from ratiometric intensity measurements. This approach is particularly advantageous for its insensitivity to optical aberrations and its compatibility with relatively high density images, which can be detrimental in PSF engineering techniques (see Supplementary Note 1). We apply this method in dense actin filament assemblies in cells, namely in the lamellipodia of motile cells and in podosomes, whose 3D organization architecture is complex and so far only accessible using electron tomography.

## Results

### Principle and data processing

The optical setup is based on a simple split of the image plane into 4 polarization channels referred to as 0°, 90°, 45° and 135°, the numerical aperture of two of which is reduced by a diaphragm in the relayed pupil plane (Fig. 1a-c) (see Materials and Methods). The diaphragm size is large enough to allow a sufficient number of photons to be detected and small enough to allow sensitivity to 3D orientations. The chosen 𝑁𝐴s lie below the critical angle aperture to avoid coupling effects with super-critical angle fluorescence (SAF) emission at the proximity to the glass coverslip interface. Typically, 𝑁𝐴_ℎ𝑖𝑔ℎ_∼1.3 (corresponding to the critical angle aperture for a glass-water interface), while the low 𝑁𝐴 value is set at about 80% of the high NA (𝑁𝐴_𝑙𝑜𝑤_∼1.1). Note that avoiding SAF also ensures that the total energy radiated by a dipole is almost identical whether it is oriented in-plane or along the axial direction^32^.

The analysis of the 4polar3D approach is based on the relative comparison of the four intensities (𝐼_0_, 𝐼_90_, 𝐼_45_, 𝐼_135_), integrated over the PSF of each single molecule, such as displayed in Fig. 1c (see Supplementary Note 1 for a description of the detailed model). The relative integrated intensities of these PSFs depend on the molecular orientation parameters, described by the molecular dipole mean 3D direction (𝜂, 𝜉), and its wobble extent 𝛿 around this direction (Fig. 1d), averaged over the integration time of the camera. We suppose that molecules are photoexcited almost isotropically (i.e. for instance using circularly polarized total internal reflection), and that their rotational diffusion is fast compared to the camera integration time. While polarized PSFs may exhibit not trivial shapes, especially for fixed and highly tilted molecules (Supplementary Fig. S1), their size is limited to the diffraction limit, in contrast to PSF engineering which expands the PSF size to at least twice this limit. This feature offers a strong advantage in situations of high detection densities. The estimation of the integrated intensities (𝐼_0_, 𝐼_90_, 𝐼_45_, 𝐼_135_) from single molecule PSFs is addressed in the section below, dedicated to the parameters’ retrieval. Supposing that these intensities are properly estimated, their dependence on the molecular angular parameters (𝜂, 𝜉, 𝛿) depicted in Figs. 1d-f (Supplementary Note 1) shows that simple combinations of (𝐼_0_, 𝐼_90_, 𝐼_45_, 𝐼_135_) provide unambiguous sensitivity to these parameters. First, the ratio between the high-NA and low-NA channels (𝐼_0_ + 𝐼_90_)/(𝐼_45_ + 𝐼_135_), is sensitive to the off-plane orientation angle 𝜂, with a higher sensitivity at low wobble angles 𝛿, as expected for fixed dipoles^31^ (Fig. 1d). Second, sensitivity to the in-plane orientation angle 𝜉 and to the wobble angle 𝛿 can be found in the respective polarization factors, i.e. 𝑃_0_ = (𝐼_0_ − 𝐼_90_)(/(𝐼_0_ + 𝐼_90_) and 𝑃_45_ = (𝐼_45_ − 𝐼_135_)/(𝐼_45_ + 𝐼_135_) ^25^. In particular, atan (𝑃_45_/𝑃_0_) is sensitive to 𝜉 (Fig. 1e), while the norm 𝑃_0_^2^ + 𝑃_45_^2^ depends on 𝛿 (Fig. 1f). These dependencies show that this scheme is capable of differentiating 3D orientation from wobble, which is not the case in a 2D polarization splitting configuration^25^. Notably, these dependencies are also robust to defocus and to the presence of unpolarised aberrations of the system, providing that numerical apertures stay under the critical angle (see Supplementary Note 1 and Suppl. Fig. S2). Note that an intrinsic property of the setup design is that 𝜉 is determined within the limit [0°-180°], and not [0°-360°], which is due to the circular geometry of the diaphragm used in the low-𝑁𝐴 channel, introducing an ambiguity between orientations symmetric with respect to the longitudinal axis 𝑧. 4polar3D thus estimates an off-plane angle η that is the amount of tilt of SMs relative to the sample plane.

The quantitative retrieval of the (η, ξ, δ) parameters from the four polarized intensities follows a simple procedure (see Supplementary Note 1). Polarized PSFs detected in the four polarized channels are linear functions of the second order moments 𝒎 of the molecular dipole vector components 𝜇_𝑖_, expressed as the time averaged quantities 𝑚_𝑖𝑗_ = 〈𝜇_𝑖_𝜇_𝑗_〉(Ω) with Ω = (η, ξ, δ) ^33^ :

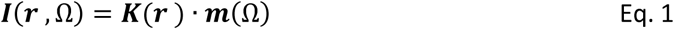

With 𝑰(𝑟, Ω) = (𝐼_0_, 𝐼_90_, 𝐼_45_, 𝐼_135_)^𝑇^the polarized PSFs, functions of 𝒓, spatial coordinate in the image plane. 𝑲(𝒓) is a 4×4 propagation matrix that depends on the detection numerical apertures as explicited in Supplementary Note 1, and 𝒎 = (𝑚_𝑥𝑥_, 𝑚_𝑦𝑦_, 𝑚_𝑧𝑧_, 𝑚_𝑥𝑦_)^𝑇^ is the vector of second order moments accessible in 4polar3D. We note that the the cross terms (𝑚_𝑥𝑧_, 𝑚_𝑦𝑧_) are absent from the estimated 𝒎 vector, as a consequence of the use of centered circular apertures as mentioned above. In 4polar3D, the total intensities 〈𝑰〉 = ∫ 𝑰(𝒓, Ω)𝑑𝒓, integrated over the whole PSF, are used to retrieve the 𝒎 components from a simple inversion operation:

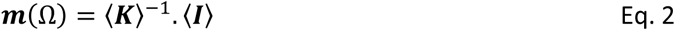

The integrated 〈𝑲〉 matrix components are determined from a calibration involving fluorescent nano-beads and a rotating polarizer (see Supplementary Note 2 and Suppl. Fig. S3), an effective way of mimicking depolarized or transverse dipoles^34^. Once 𝒎 is estimated, the retrieval of parameters Ω = (𝜂, 𝜉, 𝛿) is performed using analytical expressions of 𝒎(Ω), possibly complemented by a minimization (Supplementary Note 1)^32^.

The sensitivity of the approach to (𝜂, 𝜉, 𝛿) in the presence of noise was assessed by computing the Cramér Rao Lower Bound (CRLB) of the integrated intensities, using a typical situation of (𝑁𝐴_ℎ𝑖𝑔ℎ_ = 1.3, 𝑁𝐴_𝑙𝑜𝑤_ = 1.1) in the presence of Poisson noise and background (Supplementary Note 3). In a single-molecule regime of total intensity 5 000 photons and in the presence of 10 background photons per pixel, the estimation of the angular parameters reaches a CRLB minimal variance of about a few degrees for (𝜂, 𝜉) and less than 10° for 𝛿 (Supplementary Fig. S4). Regions of higher variances can be observed for 𝜂 ∼ 0° and 𝛿 ∼ 180°, which correspond to situations where 𝜉 is ill-defined. 𝜂 is also less precise for the extreme tilt angles (𝜂 ∼ 0°, 90°) (Fig. 1g) as expected from the 𝜂-dependence of intensity ratios displayed in Fig. 1d.

### Spatial localization and 3D orientation retrieval

The estimation of SM intensities relies on PSF shapes that are deformed by the polarization projection, especially for fixed orientations (Supplementary Fig. S1). In practice, 4polar3D employs a computationally simple procedure, in which we discard the option to run multiple-parameter fitting algorithms that would require long optimizations. After the four polarized channels have been registered, single molecules are detected in the four channels via a generalized likelihood ratio test algorithm^23,25^ (Materials and Methods). An algorithm then estimates the intensity from each PSF. We tested three options: (1) fitting the PSFs with a simplistic, albeit erroneous Gaussian function model; (2) refining this fit to a 2-dimensional Gaussian, which is still computationally light and permits to account for most deformed PSFs; (3) blindly summing pixel intensity values inside a window around each detected PSF, after subtracting the background estimated from pixels around this area. Finally, the single molecule spatial 2D localization is extracted from the center of the Gaussian estimates, or the centroid in the case of the third option. To test the precision and accuracy of intensity estimation methods, the detection, estimation and retrieval steps are performed on synthetic images generated by a Monte Carlo simulation, where theoretical PSFs are affected by binning, noise and background in different situations of image degradation conditions (Supplementary Note 4). The retrieval of (η, 𝜉, δ) is then compared to their ground truth. Despite their deformation, the sizes of polarized PSFs cover a few camera pixels as in standard SMLM detection configurations (Fig. 2a). In the absence of noise, the determination of (𝜂, 𝜉, 𝛿) is unbiased over the full range of parameter values, except for ill-defined cases mentioned above (Fig. 2b). This robustness under high signal to noise ratio (SNR) conditions has been previously exploited for the estimation of the 3D orientation of isolated single molecules under controlled orientation conditions^35^. To assess the effect of noise, background and spatial localization on the 4polar3D approach, we modeled a situation of molecules with a total intensity (sum over 4 channels) of 5 000 photons and a background of 10 photons/pixel per channel. Monte Carlo simulations show that high accuracy and precisions down to a few degrees can be reached in a range of off-plane angle and wobble (η, δ) around 20°< η < 70° and 30° < δ < 150° (Fig. 2c,d), with a negligible dependence on the in-plane angle 𝜉. In this parameter range, the intensity estimation using Gaussian fits exhibits a better performance than the box-integration estimation (Supplementary Fig. S5), which might suffer from inaccurate estimation of the surrounding background. Surprisingly, the symmetric Gaussian and rotated asymmetric Gaussian fits exhibit very similar performances, showing that a traditional detection and intensity estimation from symmetric Gaussian shapes performs well despite the deformations of polarized PSFs. In the considered parameter range (η ∈ [20° − 70°], δ ∈ [30° − 150°]) the precisions obtained from numerical simulations are close to the CRLB (Fig. 2d, Supplementary Figs. S4). At extreme values (η∼(0°, 90°), δ∼(0°, 180°)) however, a loss of performance is observed, with a degradation that depends on the number of photons as expected (Supplementary Figs. S6). This originates from the relatively lower sensitivity of the ratiometric intensity quantities to these parameters in these ranges (Fig. 1d-g).

**Fig. 2.**
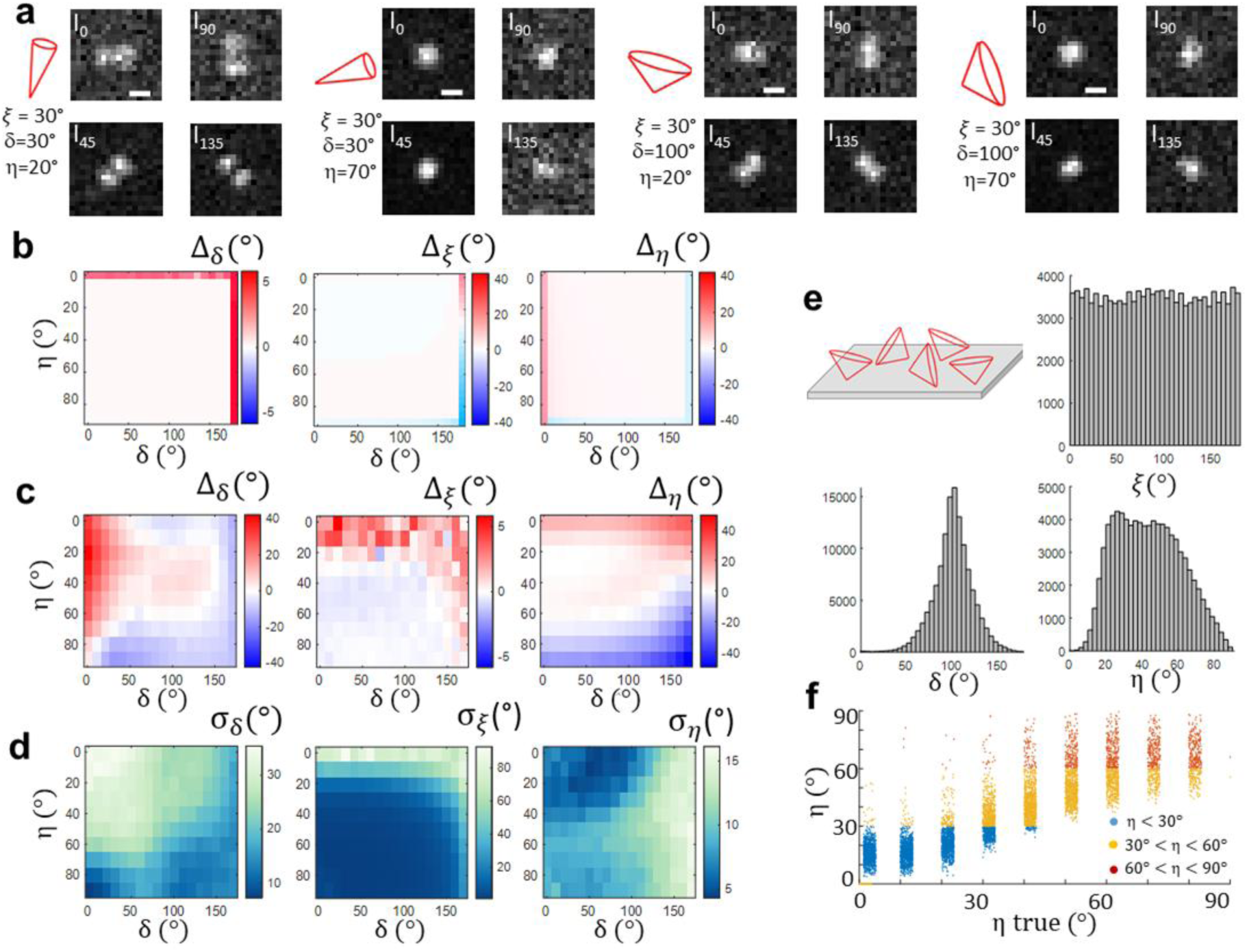
Estimation performance of 4polar3D. Conditions of Monte Carlo simulations: 10 molecules per image, 1 000 images, 5 000 photons/molecule, 10 ph/pixel background. **a** Examples of simulated PSFs (normalized to their maximum) in the presence of noise and background, with corresponding (𝜉, η, δ) values indicated on the left of each image. Scale bar: 500nm. **b** Accuracy of the parameters’ retrieval in absence of noise. **c** Accuracy of the parameters’ retrieval (Δ_δ_, Δ_ξ_, Δ_𝜂_ from left to right, values in degrees) in presence of noise, supposing the intensity estimated by a Gaussian fit. **d** Precision of the parameters’ retrieval (𝜎_δ_, 𝜎_ξ_, 𝜎_𝜂_ from left to right, values in degrees). **e** Histograms of δ, 𝜉 and η obtained from a population of random values of (𝜉, η) for δ = 100°. The population considered is schematically represented as cones of random orientations distributed on a surface. **f** Retrieved η values as functions of their ground truth value, color coded depending on the population they belong : off-plane [η < 30°], intermediate [30° < η < 60°], in-plane [η > 60°].

The consequence of the inaccuracy of the method for off-plane orientation limits is visible when simulating a collection of random orientations for a given wobble value δ (Fig. 2e). While the retrieval of δ and 𝜉 leads to expected distributions (Fig. 2e), the η histogram is cropped at its extremes η∼(0°, 90°). This effect can be mitigated when considering binned η values, opting for a data analysis that accounts for populations rather than a precise estimation of η (typically off-plane η < 30°, intermediate 30° < η < 60°, in-plane η > 60°) . Figure 2h illustrates that forming such binned populations of η, the portion of molecules belonging to the wrong population is relatively low. Finally, the different estimation methods mentioned above were tested with respect to their performance in retrieving spatial 2D localization. The simulations show that the localization precision stays below 20 nm for all used methods at a signal condition of 5 000 photons and a background of 10photons/pixel per channel, with a localization bias below 15nm (Supplementary Figs. S7). Overall, the estimation performance of 4Polar3D in orientation and spatial localization lies therefore in some parts at a lower level compared to the most efficient PSF engineering methods developed so far^1,2^, however it provides strong advantages in instrumentation and computing simplicity.

### 4polar3D SMOLM in model lipid membranes

4Polar3D was implemented experimentally using a circular excitation polarization under total internal reflection fluorescence (TIRF) illumination, in order to balance the components of the excitation polarization over the three spatial axes (see Materials and Methods). 4polar3D was first validated on lipid membranes in which the mean orientation of embedded fluorophores is predictable (Fig. 3a). Supported lipid layers (SLB) made of 1,2-dipalmitoyl-sn-glycero-3-phosphocholine (DPPC) and cholesterol mixtures (DPPC:Chol 60:40 mole%) were labeled with Nile Red (NR) (see Materials and Methods), which transiently binds to lipid membranes^11^. Within the rigid lipid environment provided by DPPC:Chol, single NR dipoles tend to orient along the axial direction of the fatty acid chains^11^, which is visible from their PSF shapes (Fig. 3b). A mean off-plane orientation of η∼30° is measured on average, with a mean wobble angle of about δ∼120° (Fig. 3c,d), which is consistent with previous findings in similar environments^11^. To reach a larger span of off-plane orientations, micrometric size spherical silica beads were then coated with DPPC/Chol lipid membranes labelled with NR, providing a well-defined wide range of 3D mean orientations similarly as in^8,11,15^ (Fig. 3e). In membrane-coated beads, both azimuthal and tilt orientations show consistent behavior with a spherical object : while azimuthal orientations ξ follow a radial trend around the sphere center at all image planes of the spheres, tilt angles η decrease from the bottom to the equator of the sphere (Fig. 3e). The η values vary between η∼30° and 80° (Fig. 3c,e), which is expected from a progressive orientation change towards the plane of the sample. The measured η range can be explained by the limits imposed by the tilt angle of NR with respect to the membrane normal direction and by the reduced η range reachable in 4polar3D. The wobble value is similar to the SLB environment, albeit with a higher heterogeneity which can be attributed to the less controlled quality of the lipid membrane on the sphere (Fig. 3c). Overall, these measurements confirm that 4polar3D SMOLM performs well in lipid membrane model systems.

**Fig. 3.**
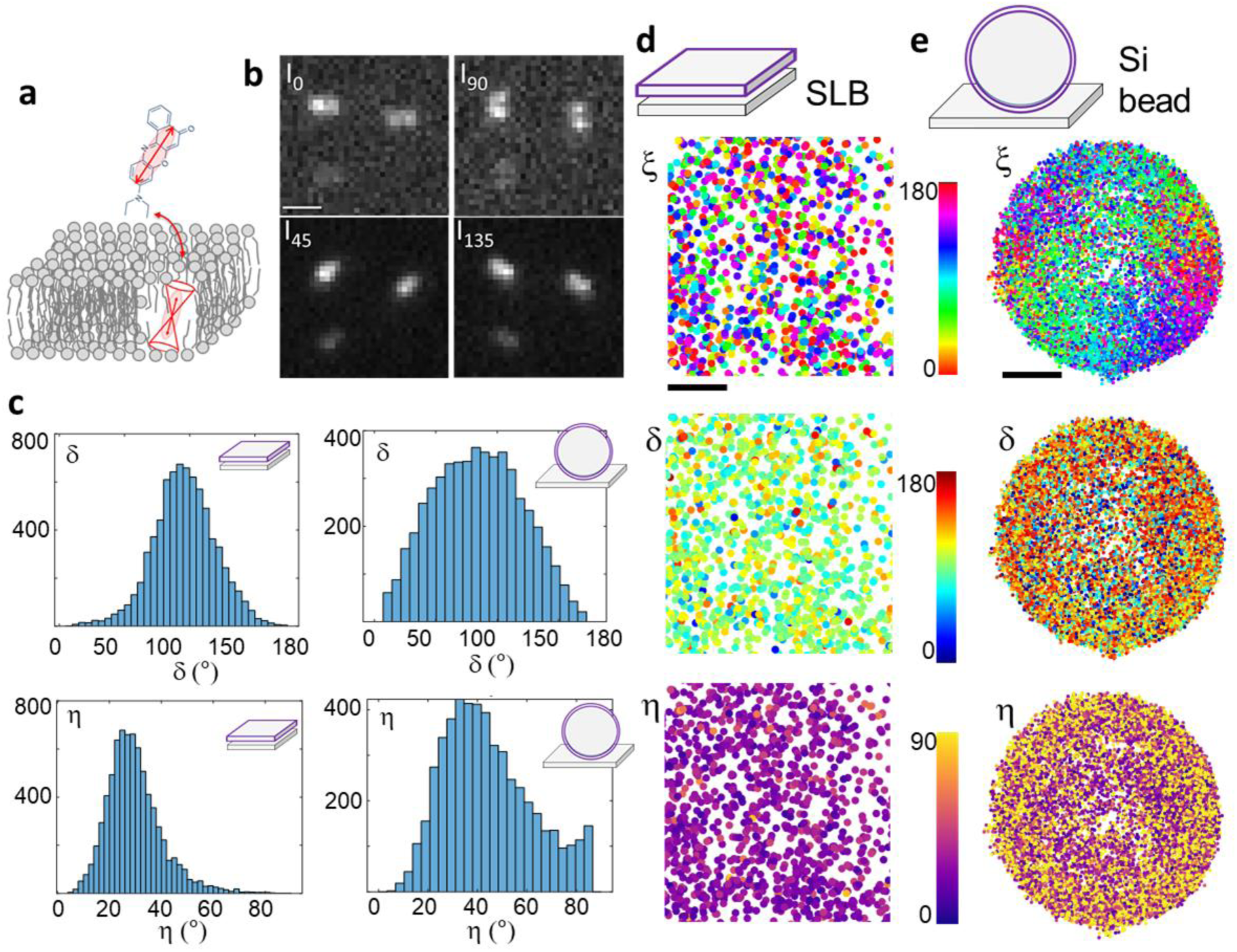
4polar3D in lipid samples. **a** Schematic representation of a lipid bilayer labelled with transiently bound Nile Red (NR) molecules. **b** 4polar3D PSFs from single NR molecules, measured in a Single Lipid Bilayer (SLB) made of DPPC(60):Chol(40) mixtures, resulting from a sum over 20 images (integration time/image 100ms). **c** Histograms of δ and η measured in SLB (left) as well as micrometric silica spheres (right) covered with lipid single bilayers of DPPC(60):Chol(40) mixtures, labelled with NR. The silica sphere data are reconstructed from different planes acquired at different tilt angles of the illumination, from the sphere bottom to close to its equator. d,e 4polar3D maps of measured angular parameters (one marker is one molecule) : ξ (top), δ (middle) and η (bottom) for (d) SLB and (e) silica micro-spheres (for which five different measured planes using different tilted illuminations, are combined in one image). Scale bars : 1μm (b), 100nm (d), 1μm (e).

### Imaging complex 3D actin filament meshworks in cells by 4polar3D SMOLM

Next, we applied the 4polar3D method to the imaging of the organization of dense actin filaments in 3D in cells. The organization of actin filaments is typically depicted as a 2D network in cells, although it is known to be fundamentally 3D in many regions of the cells, as shown by recent cryo-electron tomography works^36^.

We first focused on the lamellipodium of migrating B16 melanoma cells (Fig. 4), a thin and dense actin filament network which expands a few micrometers from the cell leading edge^37^. In this cell area, the actin filaments’ density is so large that deciphering their organization using SMLM imaging is a challenge^38^. Electron microscopy (EM) has provided evidence of 2D branched actin filament organizations at the lamellipodium edge, the branching arrangement governed by the Arp2/3 complex^37,39–41^, with some degree of heterogeneity in the way filaments arrange^42^. This network has been observed in SMOLM using the 2D version of 4polar^25^, yet direct quantification of 3D-tilted actin was not accessible. The 3D architecture of actin filaments in lamellipodia is much less studied. Recent cryo-electron tomography supported by modelling mentioned the existence of an actin network in 3D^43^, however it is unclear if the Arp2/3 branching mechanism is limited to a 2D plane. Using 4polar3D, we investigated both 3D orientation and wobble in areas where the actin filament density is high. We imaged fixed B16 cells labelled with the phalloidin conjugate AF568, which has been shown to be oriented along the actin filament direction, with a wobble of about 100° ^25^. This wobble value is non negligible, but it is expected to allow the estimation of 3D mean orientations with high precision and a reduced bias on the off-plane angle 𝜂, as seen above. Figure 4a shows a typical 4polar3D cell map of η, ξ and δ values. This whole-cell actin 4polar3D image contains 2.4 million of molecular detections obtained from a stack of 60 000 images, for which the processing time, once the detection step is done, takes about 24 seconds on a standard workstation. This time is a thousand times faster than for PSF engineering methods, which require a minimization over multiple pixels and angular parameters^26^.

**Fig. 4.**
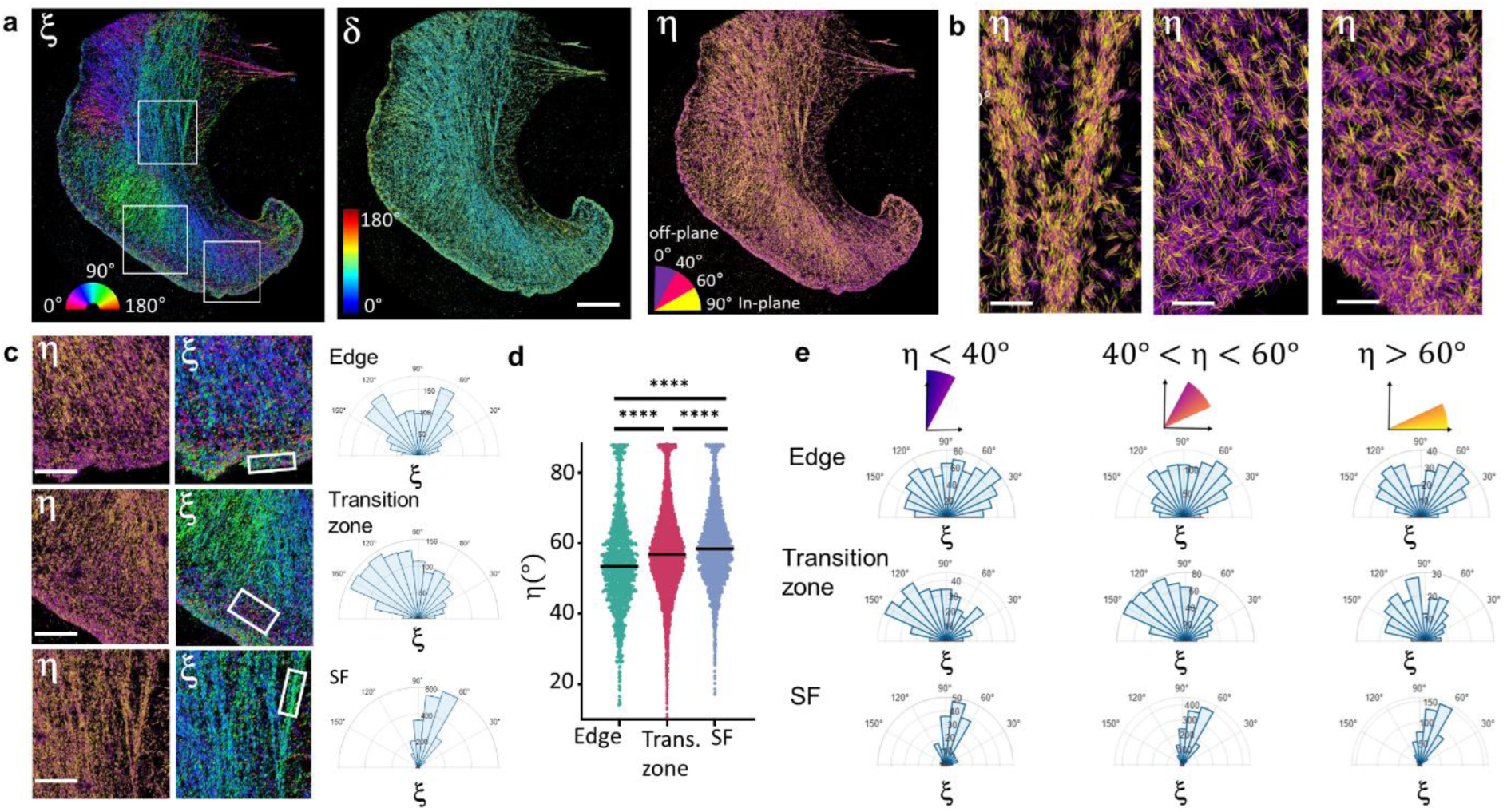
4polar3D imaging of 3D actin filaments organization in lamellipodia. **a** 4polar3D images on a fixed B16 cell labelled with phalloidin-AF568, displaying ξ (left), 𝛿 (middle) and η (right). η is shown using a binned colormap with indicated angular sectors. **b** zoomed η images of different regions (shown as white squares in (a)), illustrating different ranges of η values in stress fibers (left) and the lamellipodium region (middle, right). η is encoded as colored lines whose orientation is along ξ for each localization. **c** η and ξ zoomed images of the same cell, in three regions selected from the white rectangles in (a). For selected sub-regions (white rectangles in (c)), ξ is represented as a polar histogram (right), showing three characteristic behaviors : edge of the lamelliodium, transition zone between the lamellipodium and the cell inner region, and a stress fiber (SF). **d** η values measured in the three regions (edge, transition zone, SF) of (c) (typically, 2 500-5 000 molecules per ROI). Statistical test: unpaired t test with p values <0.0001 for all compared populations. **e** Polar histograms of ξ for given populations of η values : off-plane (η < 40°), intermediate (40° < η < 60°), in-plane (η > 60°), in the three ROIs of the cell displayed in (c). Scale bars : 5μm(a), 500nm (b), 1μm (c).

Figure 4a shows that in the lamellipodium within 1-2 μm from the cell edge, ξ spans a large range of values, which confirms the complex meshwork formed in this region. Right behind the edge, a diversity of in-plane and off-plane populations can be observed, while ventral stress fibers, located at the cell center, are clearly in-plane and directional (Figs. 4b). Distributions of ξ in zoomed regions close to the cell edge reveal the existence of bimodal distributions of actin filament orientations as previously observed^25^(Fig. 4c), which is consistent with a branched actin filament behavior promoted by Arp2/3^43^. These bimodal distributions vary along the cell contour, depicting trends with sometimes non-symmetric or broader, less well-defined distributions (Supplementary Figs. S8). Such variety of behavior has been pointed out in EM studies^43^ to be attributed to the local protruding or resting state of the cell edge^42,44^. A few micrometers away from the cell edge, in the transition zone between the lamellipodium and the lamella, ξ changes its behavior drastically, with actin filaments following the cell contour tangentially (Fig. 4c). This is in line with EM observations of long, linear bundles of actin filaments in this region^40,43,45^. In contrast, ventral stress fibers (SF) exhibit narrow and directional ξ distributions as expected (Fig. 4c). Remarkably, off-plane tilted 3D oriented actin filaments are predominantly found at the cell edge (Figs. 4a-c), which is confirmed by the measured η values in this region (Fig. 4d). We note that η does not strictly reach off-plane (η∼0°) or in-plane (η∼90°) values, due to the known bias at these extreme regions. Moreover, the anchorage of AF568 dipole to the actin filaments takes place with an offset angle of about 20° filament^25^, which means that strictly vertical filaments would lead to η∼20°. When investigating relations between in-plane and off-plane behaviors in different ROIs, we noticed that at the cell edge, the ξ bimodal distribution is more pronounced for in-plane molecules (η > 60°) than off-plane molecules (η < 40°), which depict less contrasted distributions (Fig. 4e, Supplementary Figs. S8). In contrast, distributions found in the transition zone or in SFs do not vary significantly when separating these populations. The branched actin network of the cell edge therefore seems to correspond to a predominantly in-plane population, co-existing with an off-plane population of more isotropic, less organized filaments. This behaviour has not been extensively quantified in EM studies^41,46^, even though models do not exclude 3D branched networks^47^. Specific mechanisms occuring at the very edge of the cell might be at the origin of this 3D-oriented population, where membrane proteins such as capping proteins concentrate to favor the recruitment of complexes for nucleation and elongation of new, short actin filaments^37^, ensuring crucial and dynamic remodeling^41,45^ for cell protrusion^43,44^. In contrast, the turnover rate of actin is critical at the back, a few micrometers away from the edge with long actin filaments parallel to the cell membrane^37,41,48^. The behavior of actin bundles at the transition zone and their connection to the branched population at the edge has been evidenced by EM^48^, showing the branched actin is linked to these long actin filament subsets at large distances^37^. Overall, these distinct off-plane populations might correlate with the different actin layers reported in SMLM studies^38^ or cryo-electron tomography^43^. The capacity of 4polar3D to detect and quantify the 3D organization of actin filaments in these regions emphasizes its significant added value for studies in whole cells and over large field of views. The wobble angle δ follows a rather homogeneous behavior, which shows that the mobility of the phalloidin conjugate label is overall not affected by its environment (Fig. 4b). The capability to determine δ in a 3D sensitive manner is a strong asset compared to 4polar projected in 2D^25,26^. In 2D, the measurement of δ is strongly biased when fluorophores are tilted off-plane, as confirmed by Monte Carlo simulations (Supplementary Fig. S9). This is also visible at the edge of B16 cells, where the side-by-side comparison of 4polar imaging in its 3D and 2D versions shows a significant increase of measured δ for off-plane molecules in 4polar projected in 2D (Supplementary Fig. S9).

Finally, 4polar3D was applied to the imaging of 3D actin filament organization in podosomes, which form complex actin meshworks expanding in 3D, with a lateral size of about a micrometer. Podosomes are composed of an actin-rich core able to store elastic energy and generate protrusion forces^49,50^, surrounded by an adhesion ring composed of integrins and adaptor proteins coupled to the actin core with actin lateral filaments^51^. Imaging the organization of actin filaments in this complex network is challenging due to the diffraction-limited size of podosomes, whose core is about 250-300nm wide. EM has evidenced a thin radial organization of F-actin filaments at the podosome border, with off-plane actin filaments at their core up to a thickness of about 200-400 nm^52^. This spatial architecture was later corroborated by SMLM microscopy^50,53^, although SMLM misses the information on the 3D orientation of actin filaments in these structures, leaving open questions on the co-existence of ventral in-plane and off-plane actin networks^54^ and on the organization of actin filaments between neighboring cores. Figure 5a shows a typical 4polar3D image of a fixed Human primary monocyte-derived macrophages (hMDM) cell containing a large number of podosomes, labeled by phalloidin-AF568. While the actin fibers at the cell contour have well defined orientations, podosomes contain a wide range of orientations, from which an in-plane radial behavior pointing out from the core center is perceptible (zooms, Fig. 5b). The visible long range radial behavior is in line with previous EM imaging studies, which have shown that an actin network connects closely-associated podosome cores^54^. To visualize the actin architecture inside podosomes, averaged 4polar3D images over ∼500 Podosomes (selected as ‘in’ regions in Fig. 5a and centered on their center of mass) are displayed in Fig. 5b. These images are made from typically 2 000-7 000 single molecule detections/podosome. Figure 5b shows a clear ξ radial behavior, while η decreases at the center of the podosomes, which is a signature of more off-plane filaments in the core region and gradually more in-plane filaments in the actin ring around the core. In contrast, regions between the podosomes (selected as ‘out’ in Fig. 5a and centered) display no specific ξ pattern, with dominating in-plane orientations (Fig. 5c). To quantify the radial actin behavior we introduce the radiality Δξ, difference between the in-plane angle ξ and the radial angle of each single molecule, calculated from its position relative to the center coordinate of the region (Fig. 5d, inset). Figure 5d shows that in pododomes (‘in’ regions), the portion of radial molecules (quantified by the portion for which Δξ < 10°) is maximal at a distance 𝑑 ∼400nm from the podosome center, while regions taken between the podosomes (‘out’ regions) show no specific radial pattern. The distance 𝑑 ∼400nm, which defines a typical podosome size in EM^52^, is also the one at which the in-plane actin population is the most predominant, as quantified by the 𝜂 distributions in Fig. 5e. Figure 5e clearly shows that off-plane filaments are the most abondant at the center of the podosomes, in agreement with cryo-electron tomography analyses^52^. Nevertheless, an important population of in-plane actin filaments co-exist with these off-plane filaments in the podosome core (Fig. 5b), which confirms the structural complexity of podosomes. A question that can be raised is how these different orientational populations are spatially distributed in height, over the podosome thickness. While 4polar3D does not permit to assess quantitative axial positions, the measured radius of the PSFs, when the image plane is set at the coverslip in total internal reflexion imaging conditions, permits to attribute the molecules to either ‘in-focus’ or ‘out-of focus’, keeping in mind that PSF radius also increases while η decreases (Supplementary Fig. S10). We observe that at the podosome center, in-plane/in-focus filaments (e.g. η > 60° & PSF radius < 1.4 pixels) co-exist with strongly off-plane/off-focus filaments (e.g. η < 40° & PSF radius > 2 pixels), while at the podosome border and outside podosomes, the PSFs appear to be mostly from in-focus, in-plane filaments (Fig. 5f). We note that correlations between height and off-plane orientations do not necessarily hold in the case of thick structures; indeed large ranges of PSF radii can be observed, for example, in thick stress fibers which nevertheless lie in plane (Supplementary Fig. S10). However, the maximal values of PSF radii observed in podosomes correspond to molecules lying typically off-plane and 400 nm above the coverslip, which is in line with the reported height of podosomes^50^. Our observations bring altogether new insights into the complex 3D architecture of actin networks in the podosome core, with off-plane filaments more predominantly positioned above a ventral structure, and in-plane filaments expanding radially beyond the podosome size.

**Fig. 5.**
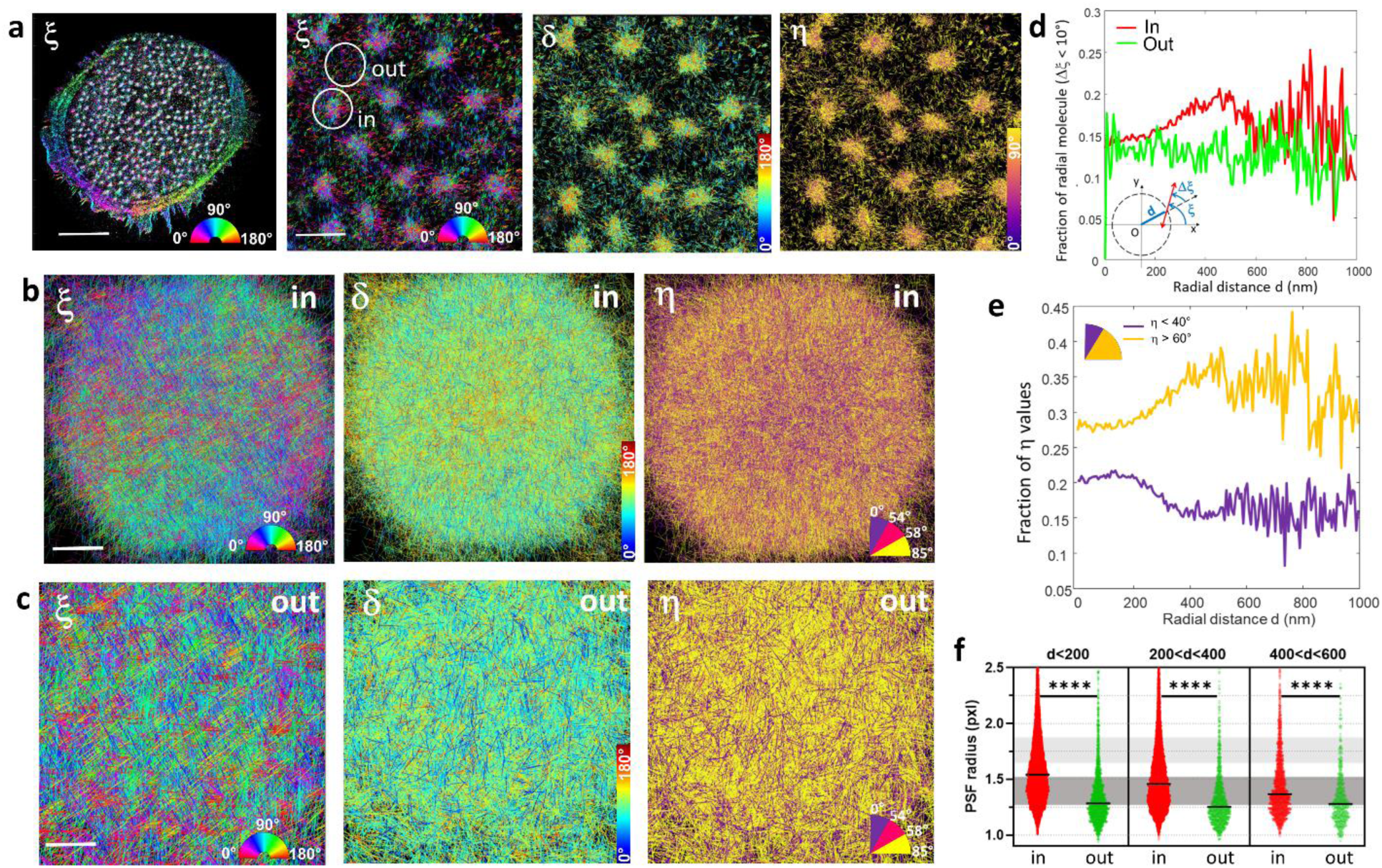
4polar3D imaging of 3D actin filaments organization in podosomes. Podosomes are imaged from Human primary monocyte-derived macrophages (hMDM) cells labelled with phalloidin-AF568. **a** In-plane orientation ξ diplayed for a whole cell (left), zoom on several podosomes showing ξ, 𝛿 and η. The circles denote the chosen regions named ‘in’ (centered on a podosome) and ‘out’ (centered on the actin cloud surrounding podosomes). **b** 4Polar3D images produced from the superposition of 504 podosomes centered on their center of mass, analyzed from 6 cells (typically, 2 000-7 000 detections per podosome ROI). Each localization is a line oriented by its in-plane orientation angle ξ, which color is coded by the corresponding parameter : ξ, 𝛿, η . η is color coded by a binned value within the displayed angular ranges. **c** Similar 4Polar3D images produced from the superposition of 807 regions of similar sizes as in (b), outside of the podosomes, analyzed from 6 cells (typically, 100-500 detections per ROI). **d** Fraction of ‘radial’ molecules over the total number of localizations (defined as deviating from less than 10° from the radial direction to the ROI center), as a function of their distance to the ROI center (values are averaged over all localizations, within 10nm distance steps), calculated over all localizations within ‘in’ (podosomes, red) and ‘out’ (actin cloud, green) ROIs displayed in (b,c). The fraction of radial molecules is defined by the proportion of the total localizations for which the radiality Δξ is between +/- 10°. The radiality Δξ is defined as the angular difference between ξ and the angle carried by the vector from the ROI center to the localization. **e** Fraction of molecules (over the total number of localizations) within ‘in’ podosome regions, oriented off-plane (η < 40^ο^) or in-plane (η > 60^ο^), as a function of their distance to the ROI center. **f** Radius of the point spread function (PSF) measured on all localizations, for different ranges of distance *d*, for both ‘in’ and ‘out’ populations within ROIs displayed in (b,c). The grey ranges of values represent the expected PSF radius of theoretical PSFs for in-plane wobbling molecules (η = 90°𝛿 = 100°, ξ random) (dark grey) and off-plane wobbling molecules (η = 0°,𝛿 = 100°, ξ random) (light grey). Scale bars = 5μm (a, full cell), 1μm (a, zooms), 200 nm (b), 100 nm (c).

## Discussion

In this work, we have demonstrated the use of an intensity-based method for single molecule 3D orientation assessment, in addition to 2D localization. This approach, named 4polar3D, relies on a simple setup implementation, that is accessible in any image-splitting microscope and requires minimal calibration steps, without the need to estimate optical aberrations in the microscope detection path. The PSFs obtained in 4polar3D are less enlarged than in PSF engineering methods, allowing the exploration of high density samples (i.e. with densely packed localizations) or dynamical situations that are challenging to PSF engineering such as single molecule tracking. The relatively simple PSF deformation also allows a straightforward estimation of their integrated intensities from SMLM-based estimation procedures, which makes signal processing both accessible and remarkably fast comparing to PSF engineering approaches. Notably, 4polar3D can be implemented in biological samples with high molecular density, which is typically the case in cells. The accuracy and precision of 4polar3D are within a few degrees for a wide range of molecular orientations in standard SNR scenarios, with limitations only at extreme orientational regimes, e.g. fixed/isotropic dipoles and totally in- or off-plane orientations. To mitigate the lower robustness of the method in these extreme regimes, we propose to implement angle binning, which is largely informative when one wants to differentiate orientational behaviors that are present in a given region of a complex sample.

We have showcased the capacity of 4polar3D to decipher the 3D organizations of actin filaments in complex actin meshworks in whole cells in lamellipodia and podosomes, over large fields of views. 4polar3D is particularly powerful for situations where statistics over a high number of regions are needed and without the need for physical sectioning or tomography, which makes this approach highly complementary to EM. Large fields of view are particularly useful for understanding transitions of orientational organizations across the cell surface at scales larger than a micrometer, and relate this organization to cell functions such as migration, protruding mechanisms and growth. Importantly, 4polar3D is fully compatible with co-localization studies of adaptor proteins or actin-binding proteins, whose nanometric scale localization is expected to strongly correlate with the structural 3D organization of actin filaments.

A significant added value of the way that 4polar3D is implemented is that it can potentially be extended to other modalities. Since 4polar3D is purely intensity-based and is robust to geometrical aberrations, its extension to multicolor imaging is straightforward and requires only wavelength-associated calibration steps to accommodate for different polarization chromatic distorsions occuring in microscope optics. It is also applicable to multiplane imaging methods, which are among the reliable approaches to achieve 3D localization over a large range of depths.

Finally, the ratiometric nature of 4polar3D makes it applicable to ensemble imaging, which is a very strong asset compared to PSF engineering approaches. In this sense, it is fully compatible with the use of genetically-encoded fluorescent-protein-based reporters for measurements of the 3D organization of actin filaments in living cells and tissues^55^. It provides a simple strategy compatible to wide field imaging, but also scanning modalities such as super resolution STED, MINFLUX, confocal and multiphoton scanning microscopy, thus potentially bringing an adequate solution to the current needs of polarized ensemble microscopy in 3D^56^.

## Materials and Methods

### 4polar3D microscope

The 4polar3D microscope is a total internal reflection fluorescence (TIRF) imaging system based on a commercial inverted microscope (Eclipse Ti2-E, Nikon) with a custom-built TIRF excitation module and polarization sensitive imaging channels. A 561 nm continuous wave laser (Coherent) is used to excite the fluorophores, which is first expanded 10x (GBE10-A, Thorlabs) before being focused (AC254-400-A-ML, Thorlabs) at the back focal plane of the 100x NA 1.45 oil immersion objective (CFI SR HP Apo TIRF 100XC Oil, Nikon). The beam expander and focusing lens are mounted on translation stage, which enables switching between epi- and TIRF illumination. Before the beam is expanded, a pair of achromatic wave plates (AQWP05M-600 and AHWP05M-600, Thorlabs) are added to correct for polarization distortions that the laser can encounter from reflections, in particular on the dichroic mirror mirror (Di01-R405/488/561/635-25×36 Semrock). This minimizes the photoselection caused by the polarization distortions of the excitation beam. For measurements using AF568 and Nile-Red fluorophores, an emission filter (FF01-609/54-25 Semrock) was used.

In the 4polar3D imaging module at the right output port of the microscope, a non-polarizing beam splitter is used to split the fluorescence emission thereby creating two optical paths each having a 1x imaging relay using pairs of 150 mm achromatic lenses (AC254-150-A-ML and AC508-150-A-ML, Thorlabs). The resulting pixel size in the object plane is 130 nm. At each of these imaging paths, an iris diaphragm is placed at their respective conjugate back focal plane to control the effective numerical apertures. Along one of the imaging paths, a half-wave plate (AHWP05M-600, Thorlabs) is placed to rotate the polarization by 45° with respect to the other path. Wollaston prisms (Quartz Wollaston Polarizer 68-820, Edmund Optics) are then placed right after each of the conjugate back focal planes producing two sets of orthogonal linearly polarized imaging paths. The imaging paths are then directed using a mirror allowing all four imaging paths to be captured by the camera (ORCA-Fusion C14440-20UP, Hamamatsu, used in ultra-quiet mode with an imaging integration time of 100-185ms.). To compensate for the polarization distorsions imposed by the dichroic mirror, a second identical dichroic mirror is mounted right after the exit port of the micrcoscope, in a vertical orientation that inverses the role of the p and s components of the fluorescence emission. This compensation dichroic is aligned to minimize the extinction for the 0/90 and 45/135 polarization channels, while observing unpolarized white light with a rotating polarizer in the emission path. Lastly, Bertrand lenses (AC254-100-A-ML and AC508-100-A-ML) are are placed on flip mounts to image the back focal plane and adjust the numerical apertures of the system. The microscope was also equipped with a three dimensional piezoelectric stage (U-780.DNS, Physik Instrumente) and the Nikon Perfect Focus System (PFS) to ensure minimal sample drift.

### Image processing

4Polar3D anaysis is run on custom MATLAB scripts. The 4 channels images are first registered using as reference a fluorescent nanobeads samples. One of the polarized channels is used as a reference for the registration, for which affine transformations are defined for the three other images (Matlab Imaging Processing toolbox). A refined registration step is then applied directly on the single molecule data in the pairing process, similarly as^25^, to refine the association of molecule’s polarized projections. For nanobeads or single molecules, we apply a detection method identical to the previously developped 4polar technique^25^, using the likelihood of detecting a single molecule spot over noise (maximum likelihood is used for the PSF peak detection within a search window of typically 11×11 pixels). This step approximates the PSFs from single molecules as symmetric Gaussian shapes. Even though this model is known to be not adapted to polarized PSFs due to their distinct orientation-sensitive shapes, especially for fixed dipoles (see Supplementary Fig. S1), this is satisfactory for the detection step, as tested in Monte Carlo simulations (see Supplementary Note 5). Molecules are coupled between two images (0-90) and (45-135) by selecting the nearest neighbor to the expected position at a vector distance from the reference image estimated from the registration step, within a distance tolerance corresponding to the localization precision. Then the four identified detections are associated. Computing the Fisher matrix and the associated Cramér Rao bound allows to estimate the theoretical lowest error expected for each PSF parameter as detailed in^57^, in particular the localization precision. This quantity is computed for all four polarization projections.

Once molecules are detected along the four channels, their parameters (2D localization, background level, PSF radius) are determined. Their intensities are estimated by PSF integration, using a PSF fit based on a Gaussian shape function (Gauss-Newton regression and minimization of a least squares analysis), a symmetric Gaussian function, or a box integration approach (see Supplementary Note 5). From the four intensities obtained along the four polarized channels, the parameters (η,ξ,δ) are estimated for each molecule using the approach described in Supplementary Note 1. Before the final image representation, drift correction is performed using the ThunderSTORM plugin on ImageJ (cross-correlation method with each bin containing 2000-5000 frames). The final 4polar3D image depicts the parameters as sticks images, using one stick per molecule detected, whose orientation relative to the horizontal axis is ξ, and whose color is either encoding ξ, η or *δ*.

### Samples preparation

#### Lamellipodia

B16-F1 cells (gift from Klemens Rottner, Technische Universität Braunschweig, Germany) were cultured in DMEM (ThermoFisher Scientific, 41966-029) supplemented with 10% fetal bovine serum (PAA Laboratories, A15-102), 100 U/mL penicillin and 100 μg/mL streptomycin (Sigma, P4333) in a humidified incubator at 37 °C and 5% CO2. B16-F1 cells were originally from ATCC (CRL-6323). H1.5 24mm diameter (170 μm ± 5 μm) high precision coverslips (Marienfeld, 0117640) were sonicated in 70% Ethanol for 5 minutes and air-dried before being coated with mouse laminin (SIGMA L2020) for 1 hour at room temperature (RT) and at a final laminin concentration of 25 μg/mL in coating buffer (50 mM Tris-HCl pH 7.5, 150 mM NaCl). B-16 cells were plated on the coated coverslips and allowed to spread overnight. To stimulate lamellipodia formation, cells were treated with aluminium fluoride for 15 min by adding AlCl_3_ and NaF solutions to a final concentration of 50uM AlCl3 and 30mM NaF. Cells were fixed for 20 min with 0.25% glutaraldehyde and 4% formaldehyde in cytoskeleton buffer (10 mM MES pH 6.1, 150 mM NaCl, 5 mM EGTA, 5 mM MgCl_2_, 5 mM glucose). The coverslips were washed twice with 1mg/ml fresh NaBH_4_ in PBS for 5 minutes each, followed by three washes with PBS for 5 minutes each. The cells were then incubated overnight with 0.5uM AF568-Phalloidin in 0.1% saponin/10% BSA in PBS at 4°C in a humidified chamber.

#### Podosomes

Monocytes from healthy subjects (HS) were provided by Etablissement Français du Sang, Toulouse, France, under contract 21/PLER/TOU/IPBS01/20130042. According to articles L12434 and R124361 of the French Public Health Code, the contract was approved by the French Ministry of Science and Technology (agreement number AC 2009921). Written informed consents were obtained from the donors before sample collection. Human peripheral blood mononuclear cells were isolated from the blood of healthy donors by centrifugation through Ficoll-Paque Plus (Cytiva), resuspended in cold phosphate buffered saline (PBS) supplemented with 2 mM EDTA, 0.5% heat-inactivated Fetal Calf Serum (FCS) at pH 7.4 and monocytes were magnetically sorted with magnetic microbeads coupled with antibodies directed against CD14 (Miltenyi Biotec #130-050-201). Monocytes were then seeded on glass coverslips at 1.5×10^6^ cells/well in six-well plates in RPMI 1640 (Gibco) without FCS. After 2h at 37°C in a humidified 5% CO2 atmosphere, the medium was replaced by RPMI containing 10% FCS and 20 ng/mL of Macrophage Colony-Stimulating Factor (M-CSF) (Miltenyi 130096489). Human primary monocyte-derived macrophages (hMDM) were harvested at day 7 using trypsin-EDTA (Fisher Scientific) and let adhere for 3h in RPMI containing 10% FCS on 1.5 H precision glass coverslips (Marienfeld). Previously, the coverslips had been cleaned by a 15 min incubation at 80°C in a RBS 35 solution (Carl Roth 9238, 1/500 diluted in milliQ water), were rinsed three times with milliQ water, let dry and sterilized at 175 °C for 120 min. For immunofluorescence, hMDM were fixed for 10 min at room temperature in a 3.7% (wt/vol) paraformaldehyde (Sigma Aldrich 158127) solution containing 0.25% glutaraldehyde (Electron Microscopy Sciences 16220) in Phosphate Buffer Saline (PBS) (Fisher Scientific). When indicated, before fixation, cells were mechanically unroofed at 37 °C using distilled water containing protease inhibitors (Thermo Scientific™ 87786) and 10 µg/mL phalloidin (Sigma-Aldrich P2141). After fixation, quenching of free aldehyde groups was performed by treatment with 50 mM ammonium chloride and 1 mg/mL NaBH4 in PBS, followed by three washes with PBS for 5 minutes each. The cells were then incubated overnight with 0.5 μM AF568-Phalloidin in 0.1% saponin/10% BSA in PBS at 4°C in a humidified chamber.

#### Nanobeads calibration sample

Fluorescent nanobeads (yellow-green Carboxylate-Modified FluoSpheres 0.1um, ThermoFisher Scientific F8803) are sonicated for 2mins, diluted in milliQ water by 10^5 times and sonicated again for 2min. Imaging spacers from Merck (GBL654004) are stuck on plasma cleaned 1.5 H coverslips. 25ul bead solution is placed in the well and allowed to dry completely. 20ul milliQ water is added and a glass slide is stuck on the spacer. The sample is sealed with nailpolish.

#### STORM imaging buffer

The final composition of the buffer for 4polar-STORM measurements was 100 mM Tris-HCl pH 8, 10% w/v glucose, 5 U/mL pyranose oxidase (POD), 400 U/mL catalase, 50 mM β-mercaptoethylamine (β-MEA), 1 mM ascorbic acid, 1 mM methyl viologen, and 2 mM cyclooctatetraene (COT). D-(+)-glucose was from Fisher Chemical (G/0500/60). POD was from Sigma (P4234-250UN), bovine liver catalase from Calbiochem/Merck Millipore (219001-5MU), β-MEA from Sigma (30070), L-ascorbic acid from Sigma (A7506), methyl viologen from Sigma (856177), and COT from Sigma (138924). Glucose was stored as a 40% w/v solution at 4 °C. POD was dissolved in GOD buffer (24 mM PIPES pH 6.8, 4 mM MgCl2, 2 mM EGTA) to yield 400 U/mL, and an equal volume of glycerol was added to yield a final 200 U/mL in 1:1 glycerol:GOD buffer; aliquots were stored at −20°C. Catalase was dissolved in GOD buffer to yield 10 mg/mL, and an equal volume of glycerol was added to yield a final 5 mg/mL (230 U/μL) of catalase in 1:1 glycerol:GOD buffer; aliquots were stored at −20°C. β-MEA was stored as ∼77 mg powder aliquots at −20 °C; right before use, an aliquot was dissolved with the appropriate amount of 360 mM HCl to yield a 1 M β-MEA solution. Ascorbic acid was always prepared right before use at 100 mM in water. Methyl viologen was stored as a 500 mM solution in water at 4 °C. COT was prepared at 200 mM in DMSO and aliquots stored at −20 °C. After mixing all components to yield the final buffer composition, the buffer was clarified by centrifugation for 2 min at 16,100 g, and the supernatant kept on ice for 15 min before use. Freshly prepared STORM buffer was typically used within a day.

#### Supported lipid bilayers

Supported lipid bilayers (SLB) were made following a solvent-assisted method^58^. Phospholipid stocks of 1,2-dipalmitoyl-sn-glycero-3-phosphocholine (DPPC) and cholesterol were purchased from Avanti Polar Lipids (Birmingham, AL, USA). In short, glass coverslips were prepared by treating them with piranha acid (3 parts of H_2_SO_4_ and 1 part of 30 wt % H_2_O_2_) for 10 min and then stored in pure water up to a month. This ensured that the glass surface is both hydrophilic and clean. Before SLB production, coverslips were exposed to oxigen plasma for 5 min and then a microfluidic chamber (Sticky-SlideVI, Ibidi) was stick on it and the channels filled with water to avoid exposure with the air. Pharmed BPT and teflon tubes were used to input isopropanol in the selected channel at 100 μL/min for 10 min followed by the input of the selected lipid mixture concentrated at 0.2 g/L at 50 μL/min for 3 min. The flow was stopped for 6 min to allow for an incubation period prior to pump Tris-NaCl buffer (10 mM Tris and 150 mM NaCl, pH=7.5) at 70μL/min for 10 min.

#### Silica beads coated with lipid bilayers

Materials: Silica beads of 3 μm diameter were purchased from Kisker Biotech GmbH & Co. KG.1,2-dipalmitoyl-sn-glycero-3-phosphocholine (DPPC) and cholesterol were purchased from Avanti Polar Lipids. Chloroform was purchased from VWR chemicals. All other reagents were purchased from Sigma-Aldrich. Methods: The method for the preparation of silica beads coated with lipid bilayers is based on previously developed protocols^26^. Preparation of silica beads: Silica bead stock at 50 mg/ml was diluted to prepare a working concentration of 2mg/ml. To prepare the 2mg/mL bead solution, the bead stock was vortexed for 10 seconds and centrifuged at 14500 rpm for 5 minutes, and the resulting pellet was resuspended in the TRIS buffer (20 mM TRIS and 50 mM KCl, pH 7.5). The vortexing and centrifugation processes were repeated 3 times. The resulting bead solution was kept in a water bath at 70 °C for 15 mins, then supplemented with 20 mM of CaCl_2_, vortexed for 30 seconds and kept in the water bath at 70 °C for further 15 minutes. Preparation of small lipid vesicles: 60 mole% DPPC and 40 mole% cholesterol were dissolved in chloroform in a glass vial. The chloroform was then evaporated under a gentle flow of nitrogen gas, resulting in a thin lipid film on the vial. The lipid film was kept overnight in a vacuum chamber to remove any excess residue of organic solvent. Next, the lipid film was rehydrated at a concentration of 1.4 mg/ml in the TRIS buffer, which was preheated to 70 °C in a water bath. The vial was sealed to prevent evaporation. The lipid suspension was incubated in a water bath at 70 °C for 30 mins. During this incubation period, the lipid suspension was vortexed every 5 mins. The hydrated lipid film was then sonicated in a bath sonicator at 70 °C for 30 minutes until the solution became completely transparent, indicating the formation of small vesicles. Preparation of beads coated with lipid bilayers: Immediately after the preparation of the small vesicles, the vesicle solution and the bead solution (at 2 mg/mL) were mixed in a volume ratio of 1:1, and vortexed to mix them well. To allow the vesicles to fuse with each other and form lipid bilayers on the beads, the vesicle-bead mixture was incubated in a water bath at 70 °C for 1 hour, while being vortexed every 10 mins during this incubation time to facilitate complete coating of the beads with lipid bilayers. The resulting beads coated with lipid bilayers were allowed to cool slowly to room temperature for 1 hour, while being vortexed every15 mins during the incubation period. To remove excess vesicles from the coated bead solution, the solution was centrifuged at 5000 rpm for 5 mins at room temperature. The supernatant was discarded, and the pellet was replenished with the TRIS buffer and vortexed well. This procedure was repeated 5 times. The bilayer coated beads were stored at 4 °C for a maximum of 2 weeks for experiments. Preparation of the SLB-coated beads with Nile Red labelling for observation: #1.5 coverslips were coated with 0.01% PLL for 20 minutes and washed thrice with Tris buffer. The SLB-coated bead solution was vortexed and mixed in Tris Buffer at 1:4 volume ratio. Nile Red is added to the solution at a final concentration of 0.75 nM. The mixture is vortexed again and added to the coverslip for observation.

#### Data availability

The 4polar3D raw image stacks are available on request from the corresponding author. A subset of data (2 GB), as well as the processed orientation/detection parameters from single-molecule data generated in this study are provided in the Source Data file, available at: https://doi.org/10.6084/m9.figshare.28890470 (will become active when the dataset is made public - private link: https://figshare.com/s/dbe846449b14b2f6078e). Processed data for example ROIs generated in this study are available for download at https://github.com/CessVala/4polar3D_SMOLM (TestData folder), with explanations provided in the README.md file.

#### Code availability

The MATLAB code used to analyse the data is available on GitHub at https://github.com/CessVala/4polar3D_SMOLM, which includes a manual and installation instructions. The Python code used for the Monte Carlo simulation is available on GitHub at https://github.com/CessVala/4polar3D_SMOLM_simulation.

## Supplementary Information

Supplementary Note 1: Theoretical principle of 4polar3D.

Supplementary Note 2: Calibration procedure and determination of the experimental 〈𝐾〉 matrix.

Supplementary Note 3: Cramér Rao Lower Bounds (CRLB) calculations.

Supplementary Note 4: 4polar3D Monte Carlo analysis.

Supplementary Fig. S1: Theoretical polarized PSFs detected in 4polar3D.

Supplementary Fig. S2: Robustness of 4polar3D to the dipole distance to coverslip.

Supplementary Fig. S3: Calibration of the 4polar3D method.

Supplementary Fig. S4: Cramér Rao Lower Bounds (CRLB) of the 4polar3D scheme.

Supplementary Fig. S5: 4polar3D Monte Carlo simulations.

Supplementary Fig. S6: Accuracy dependence on the molecule’s total intensity.

Supplementary Fig. S7: Localization accuracy and precision.

Supplementary Fig. S8: 4polar3D in the cell lamellipodia.

Supplementary Fig. S9: Estimation bias of 4polar versus 4polar3D.

Supplementary Fig. S10: 4polar3D on actin stress fibers.

## Acknowledgements

The authors warmly thank Simli Dey and Feng-Ching Tsai (Institut Curie, Paris) for the preparation of the Silica beads coated with lipid bilayers. This research has received funding from the France 2030 investment plan managed by the PIA France 2030 program IDEC Equipex+ grant (ANR-21-ESRE-0002) and the Initiative d’Excellence d’Aix-Marseille Université - A*MIDEX Institutes Cancer et Immunologie (AMX-19-IET-001) and Marseille Imaging. This work is also funded by the ANR grants 3DPolariSR (ANR-20-CE42-0003) and SETIPSS (ANR-22-CE13-0039), the « Investissements d’Avenir » program managed by the ANR (ANR-16-CONV-0001), France BioImaging national infrastructure ANR-10-INBS-04-07, Labex Cell(n)Scale ANR-11-LABX-0038 as part of the Idex PSL ANR-10-IDEX-0001-02, and from CNRS.

