## Supplementary Information for "4polar3D : Single molecule 3D orientation imaging of dense actin networks using ratiometric polarization splitting"

**Content**

Supplementary Note 1: Theoretical principle of 4polar3D.
Supplementary Note 2: Calibration procedure and determination of the experimental $\langle K \rangle$ matrix.
Supplementary Note 3: Cramér Rao Lower Bounds (CRLB) calculations.
Supplementary Note 4: 4polar3D Monte Carlo analysis.

Supplementary Fig. S1: Theoretical polarized PSFs detected in 4polar3D.
Supplementary Fig. S2: Robustness of 4polar3D to the dipole distance to coverslip.
Supplementary Fig. S3: Calibration of the 4polar3D method.
Supplementary Fig. S4: Cramér Rao Lower Bounds (CRLB) of the 4polar3D scheme.
Supplementary Fig. S5: 4polar3D Monte Carlo simulations.
Supplementary Fig. S6: Accuracy dependence on the molecule's total intensity.
Supplementary Fig. S7: Localization accuracy and precision.
Supplementary Fig. S8: 4polar3D in the cell lamellipodia.
Supplementary Fig. S9: Estimation bias of 4polar2D versus 4polar3D.
Supplementary Fig. S10: 4polar3D on actin stress fibers.

### Supplementary Notes

#### Supplementary Note 1. Theoretical principle of 4polar3D

#### 1. Polarized intensities dependencies on the angular parameters $(\boldsymbol{\eta}, \boldsymbol{\xi}, \boldsymbol{\delta})$.

The field radiated by a dipole vector $\mu$ is derived at the back focal plane of a microscope objective as[1]:

$$\tilde{E}(u) = A \ \exp[ik \ \Phi(u)] \ G(u) \ \mu \qquad \text{Eq. S1}$$

with $A$ the field amplitude. Supposing no aberrations, the phase term is written:

$$\Phi(u) = n_1 z_1 \gamma_1(u) - n_0 z_0 \gamma_0(u) + u \cdot r \qquad \text{Eq. S2}$$

$$\gamma_i(u) = \sqrt{1 - \frac{u^2}{n_i^2}} \qquad \text{Eq. S3}$$

where $n_0$ is the refractive index of the sample medium, $n_1$ the refractive index of the propagation towards the microscope (i.e. coverslip supposed to be matched with the immersion medium), $r$ is the lateral position of the dipole, $z_0$ its axial distance to the coverslip, and $z_1$ the distance between the coverslip surface and the nominal focus. $u = (u \cos\varphi, u \sin\varphi)$ is the pupil coordinate normalized so that its maximum value is the numerical aperture $NA$ of the system, the $NA$ corresponds to $u = n_0$. $G(u)$ is the $2 \times 3$ Green tensor relating each component of the dipole to the field distribution in the pupil, with[2,3] :

$$G(u) = \begin{bmatrix} g_0(u) + g_2(u)\cos 2\varphi & g_2(u)\sin 2\varphi & g_1(u)\cos\varphi \\ g_2(u)\sin 2\varphi & g_0(u) - g_2(u)\cos 2\varphi & g_1(u)\sin\varphi \end{bmatrix} \qquad \text{Eq. S4}$$

with:

$$g_{0,2}(u) = \frac{n_1\sqrt{\gamma_1(u)}}{2n_0}\left(t_p(u)\gamma_0(u) \pm t_s(u)\right)$$

$$g_1(u) = \frac{n_1\sqrt{\gamma_1(u)}}{n_0^2\gamma_0(u)} u t_p(u) \qquad \text{Eq. S5}$$

with $t_p(u)$ and $t_s(u)$ the transmission coefficients for the $p$ and $s$ polarization components through the different interfaces :

$$t_s(u) = \frac{2n_0\gamma_0(u)}{n_0\gamma_0(u) + n_1\gamma_1(u)}$$

$$t_p(u) = \frac{2n_0\gamma_0(u)}{n_1\gamma_0(u) + n_0\gamma_1(u)} \qquad \text{Eq.S6}$$

The PSF formed at the image plane of the microscope is the result of an optical Fourier transform, with a lens of focal length $f$ and magnification $m$. The field at the detector coordinate is given by :

$$E(\rho) = \frac{f}{m\lambda^2}\int \tilde{E}(u)\exp[-ik\ u.\rho/m]\ d^2u = A\ g(r,\rho)\ \mu \qquad \text{Eq.S7}$$

with $\rho$ the coordinate in the image plane. $g(r,\rho)$ is the $2 \times 3$ Green tensor at the image plane:

$$g(r,\rho) = \frac{f}{m\lambda^2} \int G(u) \exp[ik\,(n_1 z_1 \gamma_1(u) - n_0 z_0 \gamma_0(u) + u \cdot (r - \rho/m))]\; d^2u \qquad \text{Eq. S8}$$

In what follows we suppose the single dipole positioned at the center of the field of view ($r = 0$), the case of a de-centered dipole can be easily deduced and follows the same operations as below. The image of the dipole at the image plane is finally the averaged quantity $\langle\;\;\rangle$ of the field squared $|E(\rho)|^2$ over a time much longer than the fluorescence lifetime:

$$I(\rho) = \langle |E(\rho)|^2 \rangle = A^2\ \mathrm{Tr}\big[g(\rho) M g^{\dagger}(\rho)\big] \qquad \text{Eq. S9}$$

Where † denotes the transpose conjugation and $M = \langle \mu\mu^{\dagger} \rangle$ is the second moment matrix formed by the time average of squared terms $\mu\mu^{\dagger}$. A time average is equivalent to averaging over all molecular directions Ω within the angular normalized angular distribution $f(\Omega)$. The elements second moment matrix $M$ are therefore[4,5] :

$$m_{ij} = \langle \mu_i \mu_j \rangle_{\Omega} = \int f(\Omega)\, \mu_i \mu_j(\Omega) d\Omega \qquad \text{Eq. S10}$$

In what follows, we suppose that $f(\Omega)$, the molecular angular distribution explored during the integration time, is a cone shape of mean orientation (η, ξ) and wobble aperture angle δ. We suppose also that the rotation time within the cone is faster than the fluorescence lifetime of the emitter (and a fortiori faster than the integration time of the camera), and that the incident polarization is close to isotropic, i.e. does not exhibit a particular photoselected direction in the sample.

Note that due to the normalization of the distribution function $f(\Omega)$, the sum of diagonal terms $m_{ii}$ is also normalized : $m_{xx} + m_{yy} + m_{zz} = 1$. The independent quantities that can be determined here are therefore five of the six normalized components $(m_{xx}, m_{yy}, m_{zz}, m_{xy}, m_{xz}, m_{yz})$. The relation between the $m_{ij}$ components and the molecular distribution $f(\Omega)$ angular parameters is given below.

Given its linear dependence on the second moment matrix elements, the PSF can be written as a linear combination of terms from a basis of normalized PSF components[3,5,6] $I_{ij}(\rho) = \left[g^{\dagger}(\rho)\; g(\rho)\right]_{ij}$ :

$$I(\rho) = A^2 \textstyle\sum_{i,j=x,y,z} m_{ij}\, I_{ij}(\rho) \qquad \text{Eq. S11}$$

4polar3D uses integrated intensities over the PSF projected along 4 polarization channels $\sigma = x, y, p, m$ (with $p$ and $m$ along the direction 45° and 135° relative to $x$), expressed as :

$$\langle I_{\sigma}(\rho) \rangle_{\rho} = A^2 \textstyle\sum_{i,j=x,y,z} \langle I_{ij,\sigma}(\rho) \rangle_{\rho}\, m_{ij} \qquad \text{Eq. S12}$$

with $\langle\,\rangle_{\rho}$ the spatial average of the PSF expansion, similar to the sum over all pixels of the measured PSF. Examples of such polarized PSFs are given in Supplementary Fig. S1.

In what follows, we simplify the spatial averaging notation into $\langle\,\rangle_{\rho} = \langle\,\rangle$. Note that whatever the polarization projection $\sigma$, there are only four non-vanishing components in the PSF basis: $I_{xx,\sigma}(\rho)$, $I_{yy,\sigma}(\rho)$, $I_{xy,\sigma}(\rho)$, $I_{zz,\sigma}(\rho)$, with two vanishing coupling terms $I_{xz,\sigma} = I_{yz,\sigma}(\rho) = 0$. This property is valid whatever the detection condition (detection $NA$, defocus or presence of aberrations). This shows that the non-diagonal terms $m_{xz}(\Omega)$ and $m_{yz}(\Omega)$ cannot be determined via a linear polarization projection of the PSF, therefore the retrieval operation concerns only four terms of the second moment $M$ matrix. Denoting $(I_0, I_{90}, I_{45}, I_{135})$ the intensities $\langle I_\sigma \rangle$ along the $\sigma = x, y, p, m$ directions:

$$\begin{pmatrix} I_0 \\ I_{90} \\ I_{45} \\ I_{135} \end{pmatrix} = \langle K(\rho) \rangle \cdot \begin{bmatrix} m_{xx}(\Omega) \\ m_{yy}(\Omega) \\ m_{zz}(\Omega) \\ m_{xy}(\Omega) \end{bmatrix} = \langle K \rangle \cdot M(\Omega) \qquad \text{Eq. S13}$$

with $\langle K \rangle_{ij,\sigma} = \langle\, I_{ij,\sigma}(\rho) \rangle$. The $\langle K \rangle$ components correspond to crossed fields integrals $\langle K \rangle_{ij,\sigma} = \int 2\, Re\left( E_\sigma^{\mu_i}(\rho)^* E_\sigma^{\mu_j}(\rho) \right)\, d^2\rho$, which can be identically written in the Fourier plane as $\langle K \rangle_{ij,\sigma} = \int 2\, Re\left( \tilde{E}_\sigma^{\mu_i}(u)^* \tilde{E}_\sigma^{\mu_j}(u) \right)\, d^2u$. Importantly, adding a geometrical phase to $\tilde{E}(u)$ such as any geometrical aberration does not affect $\langle K \rangle_{ij}$, which is a strong advantage of the method as compared to PSF shape analysis.

Eq. S13 permits to separate the molecule angle information (i.e. mean direction orientation $(\eta, \xi)$, wobble angle $\delta$) from the microscope propagation properties (interfaces, numerical aperture, polarization projection, axial position of the dipole) contained in the propagation matrix $\langle K \rangle$. The experimental determination of $\langle K \rangle$ is described in Supplementary Note 2. A general expression of $\langle K \rangle$ can be found from the calculation of the corresponding integral elements, some of them being equal (in a perfect optical system without polarization distortions), which reduces its general expression in:

$$\langle K \rangle = \begin{bmatrix} \kappa_1 & \kappa_2 & \kappa_3 & 0 \\ \kappa_2 & \kappa_1 & \kappa_3 & 0 \\ (\kappa_1 + \kappa_2)/2 & (\kappa_1 + \kappa_2)/2 & \kappa_3 & (\kappa_1 - \kappa_2) \\ (\kappa_1 + \kappa_2)/2 & (\kappa_1 + \kappa_2)/2 & \kappa_3 & -(\kappa_1 - \kappa_2) \end{bmatrix} \qquad \text{Eq. S14}$$

$\langle K \rangle$ depends only three independent parameters only, $\kappa_1$, $\kappa_2$ and $\kappa_3$. It is also visible that the component $m_{zz}$ cannot be determined because of a degenerescence on the $\kappa_3$ value. Therefore in this form, only the 2D projection of the orientation in the sample plane is accessible, as described in[7]. The access to $m_{zz}$ can nevertheless be obtained by breaking the degenerescence on the $\kappa_3$ value by changing the numerical aperture ($NA$) of some of the detected channels, which is the core of the 4polar3D method. We suppose that the channels $(I_0, I_{90})$ are detected using a low numerical aperture $NA_{low}$, and the channels $(I_{45}, I_{135})$ detected with a high numerical aperture $NA_{high}$. Physically this can be provided by closing a diaphragm in a relay image of the pupil plane of the objective. In such situation, the $\langle K \rangle$ matrix becomes:

$$\langle K \rangle = \begin{bmatrix} \kappa_1 & \kappa_2 & \kappa_3 & 0 \\ \kappa_2 & \kappa_1 & \kappa_3 & 0 \\ \kappa_4 & \kappa_4 & \kappa_6 & \kappa_5 \\ \kappa_4 & \kappa_4 & \kappa_6 & -\kappa_5 \end{bmatrix} \qquad \text{Eq. S15}$$

Where the new coefficient $\kappa_6$ differs from $\kappa_3$ and the relation between $(\kappa_1, \kappa_2)$ and the other coefficients $(\kappa_4, \kappa_5)$ does not hold anymore. It is not surprising that modifying the detection $NA$ is sensitive to the tilt off-plane orientation angle of the dipole, since the specificity of this angle is to spread the emission energy differently between the center of the pupil plane of the objective (e.g. low

$NA$) ad its border (e.g. high $NA$). Below we give a typical $\langle K \rangle$ matrix expression, obtained for molecules at the coverslip surface (i.e. $z_0 = z_1 = 0$, with $n_2 = 1.33$ ; $n_1 = 1.51$). We suppose $NA_{high} = 1.3$ (i.e. super critical angular fluorescence (SAF) is not included) and $NA_{low} = 1.1$:

$$\langle K \rangle = \begin{bmatrix} 0.353 & 0.003 & 0.075 & 0.000 \\ 0.003 & 0.353 & 0.075 & 0.000 \\ 0.322 & 0.322 & 0.307 & 0.584 \\ 0.322 & 0.322 & 0.307 & -0.584 \end{bmatrix} \qquad \text{Eq. S16}$$

A typical experimentally measured $\langle K \rangle$ matrix, using the procedure described in Supplementary Note 2, is given below. It can be visible that the presence of slight polarization distorsions remove some equalities between the $\langle K \rangle$ components:

$$\langle K \rangle_{exp} = \begin{bmatrix} 0.399 & -0.001 & 0.136 & 0.005 \\ 0.008 & 0.377 & 0.105 & 0.001 \\ 0.298 & 0.310 & 0.267 & 0.493 \\ 0.293 & 0.311 & 0.287 & -0.489 \end{bmatrix} \qquad \text{Eq. S17}$$

**2. Retrieval of the angular parameters $(\boldsymbol{\eta}, \boldsymbol{\xi}, \boldsymbol{\delta})$ from four polarized intensities.**

Eq. S13 shows that the four $(m_{xx}, m_{yy}, m_{zz}, m_{xy})$ second moment components can be, in principle, directly determined from the measurement of four integrated intensities $(I_0, I_{90}, I_{45}, I_{135})$ though a matrix inversion operation:

$$\begin{pmatrix} m_{xx} \\ m_{yy} \\ m_{zz} \\ m_{xy} \end{pmatrix} = \langle K \rangle^{-1} \begin{pmatrix} I_0 \\ I_{90} \\ I_{45} \\ I_{135} \end{pmatrix} \qquad \text{Eq. S18}$$

Where $\langle K \rangle^{-1}$ is the pseudoinverse of the non-square $\langle K \rangle$ matrix. The angular parameters $(\eta, \xi, \delta)$ can then be determined from the $m_{ij}$ elements, using Eq. S10. Supposing a fast rotation of the emitter dipole (i.e. with a rotation time faster than its fluorescence lifetime) within a wobble cone of aperture $\delta$ and mean orientation $(\eta, \xi)$, the expression of the moments elements are[5]:

$$\begin{aligned} m_{xx} &= \left(1 - 3\lambda(\delta)\right) \sin^2\eta \cos^2\xi + \lambda(\delta) \\ m_{yy} &= \left(1 - 3\lambda(\delta)\right) \sin^2\eta \sin^2\xi + \lambda(\delta) \\ m_{zz} &= \left(1 - 3\lambda(\delta)\right) \cos^2\eta + \lambda(\delta) \\ m_{xy} &= \left(1 - 3\lambda(\delta)\right) \sin^2\eta \cos\xi \sin\xi \\ \text{with } \lambda(\delta) &= \tfrac{1}{6}\left(1 - \cos\left(\tfrac{\delta}{2}\right)\right)\left(2 + \cos\left(\tfrac{\delta}{2}\right)\right) \end{aligned} \qquad \text{Eq. S19}$$

Note that slower motions can be treated using more complex expressions accounting for the incident polarization[8,9].

To deduce the angular parameters $(\eta, \xi, \delta)$ we introduce reduced factors:

$$P_{uv} = \frac{2m_{xy}}{m_{xx} + m_{yy} + m_{zz}} = (1 - 3\,\lambda)\ \sin^2\eta \sin 2\xi$$
$$P_{xy} = \frac{m_{xx} - m_{yy}}{m_{xx} + m_{yy} + m_{zz}} = (1 - 3\,\lambda)\ \sin^2\eta \cos 2\xi$$
$$P_{zz} = \frac{m_{zz}}{m_{xx} + m_{yy} + m_{zz}} = (1 - 3\,\lambda)\ \cos^2\eta + \lambda$$

Eq. S20

In practice, a normalization is first performed such that $m_{xx} + m_{yy} + m_{zz} = 1$ and the searched parameters are deduced from:

$$\xi = \frac{1}{2}\tan\frac{P_{uv}}{P_{xy}}$$
$$\delta = 2\ \mathrm{acos}\left[\frac{-1 + \sqrt{9 - 24\lambda}}{2}\right]$$
$$\eta = \mathrm{acos}\left[\sqrt{\frac{P_{zz} - \lambda}{1 - 3\lambda}}\right]$$

Eq. S21

with:

$$\lambda = \frac{1}{2}\left[1 - \left(P_{zz} + \sqrt{P_{uv}^2 + P_{xy}^2}\right)\right]$$

Eq. S22

The advantage of this approach is that the determination of the $(\eta, \xi, \delta)$ from the measured four intensities $(I_0, I_{90}, I_{45}, I_{135})$ is non-ambiguous and very fast. Note that whenever the experimentally obtained dipole moments do not match the model and fall outside of the required domain of realistic $(m_{xx}, m_{yy}, m_{zz}, m_{xy})$ values (due to noise essentially), there is no physical solutions found. In practice, the closest $m_{xy}$ ensuring the condition $\lceil m_{xy} \rceil \leq \sqrt{m_{xx} m_{yy}}$ can be found, to ensure that the retrieved angles are physically meaningful. Another approach is least squares minimization, which can be used to deduce the most probable orientation $(\eta, \xi)$ and wobbling angle $\delta$ that fit the measured intensities, minimizing:

$$\min_{\eta,\xi,\delta} \sum \left[\langle K \rangle^{-1} \begin{pmatrix} I_0 \\ I_{90} \\ I_{45} \\ I_{135} \end{pmatrix} - \begin{pmatrix} m_{xx}(\eta, \xi, \delta) \\ m_{yy}(\eta, \xi, \delta) \\ m_{zz}(\eta, \xi, \delta) \\ m_{xy}(\eta, \xi, \delta) \end{pmatrix}\right]^2$$

Eq. S23

Where the sum $\sum$ is performed over the four polarization channels elements.

#### 3. Comparison between 4polar3D and the 2D version of 4polar.

In a 4polar2D experiment, the two $NA$'s of the polarized channels are equal, which leads to no access of the off-plane orientation[7,10]. Therefore $m_{zz}$ is ignored and the relevant moments quantities are:

$$\begin{pmatrix} m_{xx} \\ m_{yy} \\ m_{xy} \end{pmatrix} = \langle K \rangle_{2D}^{-1} \cdot \begin{pmatrix} I_0 \\ I_{90} \\ I_{45} \\ I_{135} \end{pmatrix} \qquad \text{Eq. S24}$$

Where $\langle K \rangle_{2D}$ matrix is a subset of the $\langle K \rangle$ matrix, missing the 3rd column.

To deduce the angular parameters from the moments, we introduce the relevant factors, deduced from Eq. S20 with $\eta = \pi/2$ due to the reduced 2D geometry of the measurement :

$$P_{uv-2D} = \frac{2m_{xy}}{m_{xx} + m_{yy}} = \frac{(1-3\lambda)}{(1-\lambda)} \sin 2\xi$$
$$P_{xy-2D} = \frac{m_{xx} - m_{yy}}{m_{xx} + m_{yy}} = \frac{(1-3\lambda)}{(1-\lambda)} \cos 2\xi$$

Eq. S25

The in-plane angle $\xi_{2D}$ (obtained from this 2D scheme) is directly deduced from the expression:

$$\tan 2\xi_{2D} = \frac{P_{uv-2D}}{P_{xy-2D}} = \frac{2m_{xy}}{m_{xx} - m_{yy}} \qquad \text{Eq. S26}$$

To measure $\delta_{2D}$, which is a projection of $\delta$ in the sample plane, we use :

$$P_{2D} = \sqrt{P_{xy-2D}{}^2 + P_{uv-2D}{}^2} = \sqrt{\frac{(m_{xx}-m_{yy})^2 + 4m_{xy}{}^2}{(m_{xx}+m_{yy})^2}} = \frac{[1-3\lambda]}{[1-\lambda]} \qquad \text{Eq. S27}$$

Where $\lambda$ is now obtained supposing the cone distribution lies in the sample plane ($\eta = \pi/2$). Using its estimated value $\lambda_{2D} = \frac{1-P_{2D}}{3-P_{2D}}$, we can then deduced :

$$\delta_{2D} = 2 \operatorname{acos}\left[\frac{-1 + \sqrt{9 - 24\lambda_{2D}}}{2}\right]$$
$$\xi_{2D} = \frac{1}{2} \tan \frac{P_{uv-2D}}{P_{xy-2D}} \qquad \text{Eq. S28}$$

This scheme was used in recent works for 2D orientation retrieval[7,10]. Supplementary Fig. S9 shows that the 2D retrieved solution induces a strong bias on $\delta$, as expected from its projection in the sample plane, but does not induce a visible bias on $\xi$.

**Supplementary Note 2. Calibration procedure and determination of the experimental $\langle K \rangle$ matrix.**

Following Eq. S13, each column of $\langle K \rangle$ dictates how a second order moment element $m_{ij}$ is imaged onto each of the four linearly polarized channels. In other words, to determine the matrix elements $\langle K \rangle$ it is sufficient to use intensity measurements from dipoles oriented in plane ($\eta$ = 90° at $\xi$ = 0°, 90°, 45° or 135°) and completely out of plane ($\eta$ = 0°) and solve the matrix multiplication since the second order dipole moments $m_{xx}$, $m_{yy}$, $m_{xy}$, and $m_{zz}$ are well defined functions of $\eta, \xi$ and $\delta$. Ideally, immobile fluorophores with known orientation would be used for this purpose, unfortunately fabricating such samples is not straightforward. In place of these ideal calibration samples, and to provide easily reproducible calibration protocols, fluorescent nanobeads with a linear polarizer are used. Fluorescent beads (20 nm size) deposited on a coverslip (see Materials and Methods) are imaged using a linear polarizer positioned at the back focal plane of the 4polar3D microscope. Fluorescent nanobeads are considered as depolarized emitters, due to the random orientation and high density of their embedded fluorescent emitters. This allows the calibration to account for possible polarization distortions brought by the optics inside the microscope, in particular the dichroic mirror (see Supplementary Fig. S3). To compensate for polarization distortions, a second identical dichroic is placed at the output of the microscope body, oriented in a way that the roles of the $s$ and $p$ polarization components are exchanged and therefore compensate their phase and amplitude in the propagation process. A set of intensity measurements $I_\sigma(\alpha)$ are performed at each polarizer orientation $\alpha$ with respect to the horizontal (0°) direction, which can be written :

$$I_{beads,\sigma}(\alpha) \propto \int \left[ \left| \left( \tilde{E}^{\mu_x}(u) + \tilde{E}^{\mu_y}(u) + \tilde{E}^{\mu_z}(u) \right) \right|^2 p(\alpha) \cdot e_\sigma \right] d^2u \qquad \text{Eq. S29}$$

Where $\tilde{E}(u)$, $p(\alpha)$ and $e_\sigma$ are vectors : $\tilde{E}^{\mu_i}(u)$ is the field radiated by the dipole component $\mu_i$, function of the pupil plane coordinate $u$, $p(\alpha)$ is the linear polarizer direction at the back focal plane and $e_\sigma$ characterizes the projection on the polarized detection channel. It should be emphasized that the intensity obtained from combining an isotropic emitter (a fluorescent nanobead) and a linear polarizer is slightly different from the emission from an oriented dipole modelled above[11]. For a dipole lying in the sample plane at an angle $\alpha$ with respect to $x$, the dipole intensity can be written as:

$$I_{dipole,\sigma}(\alpha) = \iint_0^{NA_i} \left| \left( \tilde{E}^{\mu_x}(u,\alpha) + \tilde{E}^{\mu_y}(u,\alpha) + \tilde{E}^{\mu_z}(u,\alpha) \right) \cdot e_\sigma \right|^2 \qquad \text{Eq. S30}$$

To experimentally determine the propagation matrix $\langle K \rangle$, $I_{beads,\sigma}(\alpha)$ is therefore first corrected to approximate $I_{dipole,\sigma}(\alpha)$ and thus mimick at best individual radiating horizontal dipoles along $\alpha$. For this, the intensity measurements from the fluorescent beads $I_{beads,i}(\alpha)$ can be rewritten as a simple function:

$$I_i(\alpha) \propto A_{0,i} + A_{2,i} \cos 2\alpha + B_{2,i} \sin 2\alpha \qquad \text{Eq. S31}$$

Where $i = (0,90,45,135)$. Experimentally, intensities are first normalized to the sum of orthogonal channels ((0°, 90°) on one side and (45°, 135°) on the other side). The factors $A_{0,i}$ , $A_{2,i}$ , and $B_{2,i}$ are then obtained by a direct circular projection. Note that the amplitude components $A_{0,i}$ allows a direct experimental verification of the back focal plane reduction factor $\mathrm{NA}_{low}/\mathrm{NA}_{high}$. Correction factors are then brought directly on the coefficients $A_{0,i}^{beads}$, $A_{2,i}^{beads}$, and $B_{2,i}^{beads}$ to mimic dipoles of corresponding orientations $\alpha$. To calculate these correction factors, we compute theoretical beads and dipole responses, using the known back focal plane reduction factor. Theoretical fluorescent beads and rotating dipole intensities can be calculated with their corresponding coefficients :

$$I_{dipole,i}(\alpha) = A_{0,i}^{dipole} + A_{2,i}^{dipole}\cos 2\alpha + B_{2,i}^{dipole}\sin 2\alpha$$

$$I_{beads,i}(\alpha) = A_{0,i}^{beads} + A_{2,i}^{beads}\cos 2\alpha + B_{2,i}^{beads}\sin 2\alpha \qquad \text{Eq. S32}$$

From these coefficients, correction factors are calculated which can be used on the experimental coefficients $A_{0,i}^{expt}$, $A_{2,i}^{expt}$, and $B_{2,i}^{expt}$ so that they approximate the emission of oriented dipoles.

$$A_{0,i}^{corr} = A_{0,i}^{expt}\frac{A_{0,i}^{dipole}}{A_{0,i}^{beads}},\quad A_{2,i}^{corr} = A_{2,i}^{expt}\frac{A_{2,i}^{dipole}}{A_{2,i}^{beads}},\quad B_{2,i}^{corr} = B_{2,i}^{expt}\frac{B_{2,i}^{dipole}}{B_{2,i}^{beads}}$$

$$I_{corr,i}(\alpha) = A_{0,i}^{corr} + A_{2,i}^{corr}\cos 2\alpha + B_{2,i}^{corr}\sin 2\alpha \qquad \text{Eq. S33}$$

Finally this intensity expression can be used to determine the in-plane coefficients of the propagation matrix $\langle K\rangle$ ($K_{xx,i}$, $K_{yy,i}$, and $K_{xy,i}$) by expressing the second order dipole moments as:

$$\begin{aligned} m_{xx}(\alpha) &= \cos^2\alpha \\ m_{yy}(\alpha) &= \sin^2\alpha \\ m_{zz} &= 0 \\ m_{xy}(\alpha) &= \cos\alpha\,\sin\alpha \end{aligned} \qquad \text{Eq. S34}$$

Solving Eq. S34 gives finally:

$$\begin{aligned} K_{xx,i} &= A_{0,i}^{corr} + A_{2,i}^{corr} \\ K_{yy,i} &= A_{0,i}^{corr} - A_{2,i}^{corr} \\ K_{xy,i} &= 2B_{2,i}^{corr} \end{aligned} \qquad \text{Eq. S35}$$

The last remaining column of the $\langle K\rangle$ matrix, $K_{zz,i}$, defines the sensitivity of 4polar3D imaging out of plane oriented dipoles ($m_{zz}$) . Since it is challenging to fix the out of plane components of a calibration sample a different strategy is used to determine $K_{zz,i}$. Following Eq. S14, the intensities measured by 4polar3D are given by :

$$I_i = K_{xx,i}m_{xx} + K_{yy,i}m_{yy} + K_{zz,i}m_{zz} + K_{xy,i}m_{xy} \qquad \text{Eq. S36}$$

such that

$$K_{zz,i} = \frac{I_i - K_{xx,i}m_{xx} + K_{yy,i}m_{yy} + K_{xy,i}m_{xy}}{m_{zz}} \qquad \text{Eq. S37}$$

Now consider a perfectly isotropic incoherent emitter or a completely isotropic dipole ($\delta$ = 180°) which is in fact the case for fluorescent beads with no linear polarizer. The second order moments of such emitters are $m_{xx}^{iso} = m_{yy}^{iso} = m_{zz}^{iso} = \frac{1}{3}$ and $m_{xy}^{iso}$= 0, which then reduces the expression of $K_{zz,i}$ to : $K_{zz,i} = 3I_i - K_{xx,i} - K_{yy,i}$. Once the in-plane components of the propagation matrix $\langle K\rangle$ have been calibrated, $K_{zz,i}$ can thus be directly determined by measuring isotropic fluorescent beads and solving the above equation.

**Supplementary Note 3. Cramér Rao Lower Bound (CRLB) calculations**

The Cramér-Rao lower bound (CRLB) is the lowest variance that can be obtained in the retrieval of a parameter, given a measured quantity that follows a known statistical distribution. It is defined as the inverse of the diagonal elements of the Fisher information matrix F, which expression is:

$$[\mathrm{F}(p)]_{mn} = -E\left[\frac{\partial^2}{\partial p_m \partial p_n}\ell(p|y)\right] \qquad \text{Eq. S38}$$

where $p$ is the vector containing the set parameters to be estimated ($p^T = (\eta, \xi, \delta)$), $y$ is the outcome of a measurement, and $\ell(p|y)$ is the log-likelihood function, i.e. the logarithm of the probability to obtain the set of parameters $p$, given a measurement $y$. In the context of 4polar3D, and $y$ is the measure of the total intensity projected onto the 4polarized channels. The CRLB on the variance of the parameter $p_n$ can be calculated as :

$$\sigma^2(p_n) \geq [\mathrm{F}(p)^{-1}]_{mn} \qquad \text{Eq. S39}$$

To calculate the Fisher matrix, we suppose that the integrated intensities of the PSF measured on the camera, $I^{meas} = (I_0^{meas}, I_{90}^{meas}, I_{45}^{meas}, I_{135}^{meas})$ follows a Poisson distribution centered on the expected value of the intensity, $I^{exp}$. The likelihood of measuring the intesnity value $I^{meas}$ is therefore

$$\mathcal{L}(I^{meas}|I^{exp}) = \prod_{i=1}^{4}\left(I_i^{exp}\right)^{I_i^{meas}} \cdot \frac{e^{-I_i^{exp}}}{I_i^{meas}!} \qquad \text{Eq. S40}$$

which natural logarighm becomes:

$$\ell(I^{meas}|I^{exp}) = \sum_{i=1}^{4} I_i^{meas}.\ln\left(I_i^{exp}\right) - I_i^{exp} - \ln(I_i^{meas}!) \qquad \text{Eq. S41}$$

therefore

$$\frac{\partial^2}{\partial p_m \partial p_n}\ell(I^{exp}|I^{meas}) = \sum_{i=1}^{4} -\frac{I_i^{meas}}{\left(I_i^{exp}\right)^2}.\frac{\partial I_i^{exp}}{\partial p_m}.\frac{\partial I_i^{exp}}{\partial p_n} + \left(\frac{I_i^{meas}}{I_i^{exp}} - 1\right).\frac{\partial^2 I_i^{exp}}{\partial p_m \partial p_n} \qquad \text{Eq. S42}$$

The expectation value of this expression for $I_i^{exp} = I_i^{meas}$ becomes:

$$[\mathrm{F}(p)]_{mn} = \sum_{i=1}^{4} -\frac{1}{I_i^{exp}}.\frac{\partial I_i^{exp}}{\partial p_m}.\frac{\partial I_i^{exp}}{\partial p_n} \qquad \text{Eq. S43}$$

From the construction of the 4polarized channels:

$$I_i^{exp} = \sum_{k,l=x,y,z} N\,\langle K\rangle_{ikl}.m_{kl} + b_i \qquad \text{Eq. S44}$$

with $N$ the total number of photon emitted by the molecule and $b_i$ the background level present in channel $i$. Therefore:

$$[\mathrm{F}(p)]_{mn} = N\sum_{i=1}^{4}\sum_{k,l=x,y,z}\frac{(\langle K\rangle_{ikl})^2}{\langle K\rangle_{ikl}.m_{kl}+b_i/N}.\frac{\partial m_{kl}(p)}{\partial p_m}.\frac{\partial m_{kl}(p)}{\partial p_n} \qquad \text{Eq. S45}$$

With $p^T = (\eta, \xi, \delta)$. Typical CRLB values are shown for the parameters $(\eta, \xi, \delta)$ in Supplementary Fig. S4.

**Supplementary Note 4. 4polar3D Monte Carlo analysis.**

For Monte Carlo simulations, custom scripts were written in Python using the user-friendly interface of Jupyter notebook. The 4polar 3D SMOLM simulation software is available at https://github.com/CessVala/4polar_3D_SMOLM_simulation.

Simulations were implemented to study the effect of noise and background on the retrieved 4polar3D channels intensity, using the different estimation strategies (symmetric Gaussian, rotated asymmetric Gaussian, and pixel integration). The starting parameters are the total intensity (2 500 photons, 5 000 photons, and 10 000 photons), background pixel value (10 photons/pixel), and the angular $(\eta, \xi, \delta)$ parameters. Synthetic PSF images were created using a custom MATLAB script, following the model described in Supplementary Note 1. The molecules are supposed to lie at the coverslip surface (i.e. $z_0 = z_1 = 0$), $n_2 = 1.33$ ; $n_1 = 1.51$, $NA_{high} = 1.33$ (below critical angle) and $NA_{low} = 1.16$. Note that no variation is expected at higher distances to the coverslip surface or defocus, due to the use of a sub-critical angle $NA$ regime (see Supplementary Fig. S2). Images containing PSFs without noise were first generated to which camera noise and background were added, as described previously[7]. In short, the noise statistics of the camera follows a model determined empirically by plotting the standard deviation of measured intensities at growing intensity values. For each polarized channel. $I_i$ is first normalized to the sum of all four channels intensities, $I_{tot} = \sum_i I_i$, before noise and background addition. The synthetic data consist of 100 images containing each 10 molecules, resulting in 1 000 molecules per condition, projected on the 4 polarization channels.

For the analysis, single molecules' local maxima are first determined from synthetic data. These maxima are estimated in each polarized channel using the "peak_local_max" function from the "scikit-image" python library. The main accessible parameters for this estimation are "min_distance" and "threshold intensity", which determine candidate molecules. Based on these candidate molecules, intensity estimation is performed based on three approaches (symmetric Gaussian, rotated asymmetric Gaussian, pixel integration centered on the center coordinate determined by a symmetric Gaussian fit). For Gaussian fits, the "curve_fit" function is used from the "scipy" python library to estimate the molecule localization, and its intensity is estimated as the integrated signal under the fitted PSF. For the pixel integration method, each PSF intensity is estimated as the sum of the pixels inside a 17x17 pixels window minus the median of the external pixels multiplied by the number of pixels. An adaptation of the window to the PSF size (obtained from a Gaussian fit) can also be implemented. Using parallel processing, the program can process multiple images simultaneously. After the intensity estimation is complete, registration is performed first between the quadrants $(I_0, I_{90})$ and $(I_{45}, I_{135})$, separately then one extra registration step is performed between $(I_0, I_{45})$. This allows for the retrieval of the 4 polarized projections corresponding to each simulated molecule. Then the parameters $(\eta, \xi, \delta)$ are estimated for each molecule, as described in Supplementary Note 1. The accuracy obtained for these parameters is calculated by subtraction to the ground truth, while the precision is estimated by the standard deviation obtained over the 1 000 realizations per angle conditions. The results are shown in Supplementary Figs. S5-S7 for several photon number conditions and comparing different fitting methods.

At last, the spatial localization precision is estimated later using the method of[12]. Since the expected localization of each simulated particle is known, the localization precision is defined as the standard deviation of the distribution of the 1 000 simulated localizations, assuming a Gaussian distribution, obtained along the $x$ and $y$ axes. The reported localization precisions are the best average between $x$ and $y$ obtained among the 4 channels. The accuracy in localization is estimated using the expected center of an unpolarized molecule. Typically for 1 000 detections in the presence of noise, the average processing time is about 10 sec for symmetric Gaussian fitting and box integration estimation, and a few seconds more for asymmetric Gaussian fitting.

**Supplementary Figures**

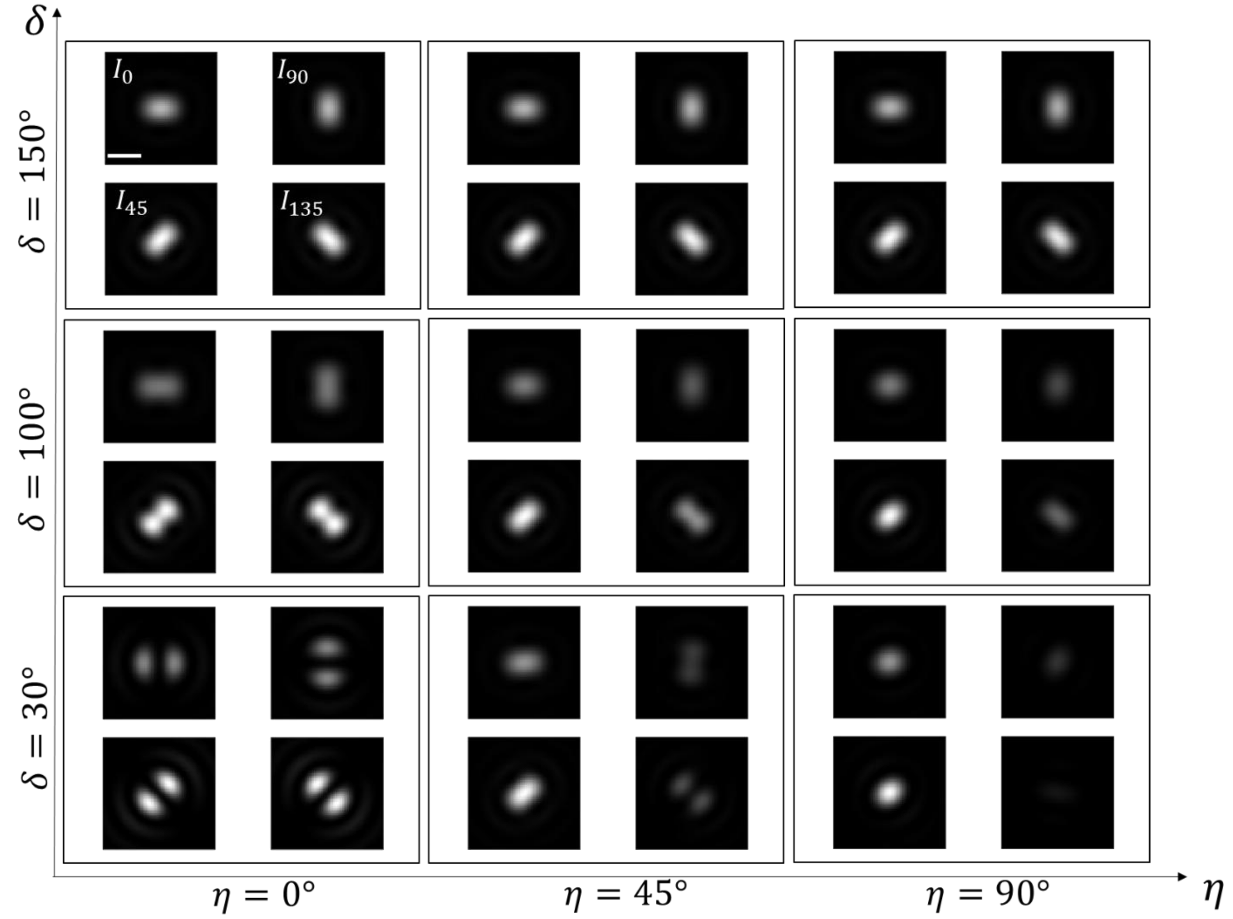


**Supplementary Fig. S1**. **Theoretical polarized PSFs detected in 4polar3D**, for various $(\eta, \delta)$ parameters and a fixed in-plane orientation $\xi = 30°$. Other $\xi$ angles lead to similar PSF shapes with a variation of the relative magnitude of the intensities $(I_0, I_{90}, I_{45}, I_{135})$. The PSF images are normalized with respect to the maximum of the total intensity $I_0 + I_{90} + I_{45} + I_{135}$. Conditions used for the simulations: $n_0 = 1.33$, $n_1 = 1.515$, $NA_{low} = 1.1$, $NA_{high} = 1.3$, distance to the coverslip $z_0 = 0$, defocus $z_1 = 0$. Scale bar : 800nm.

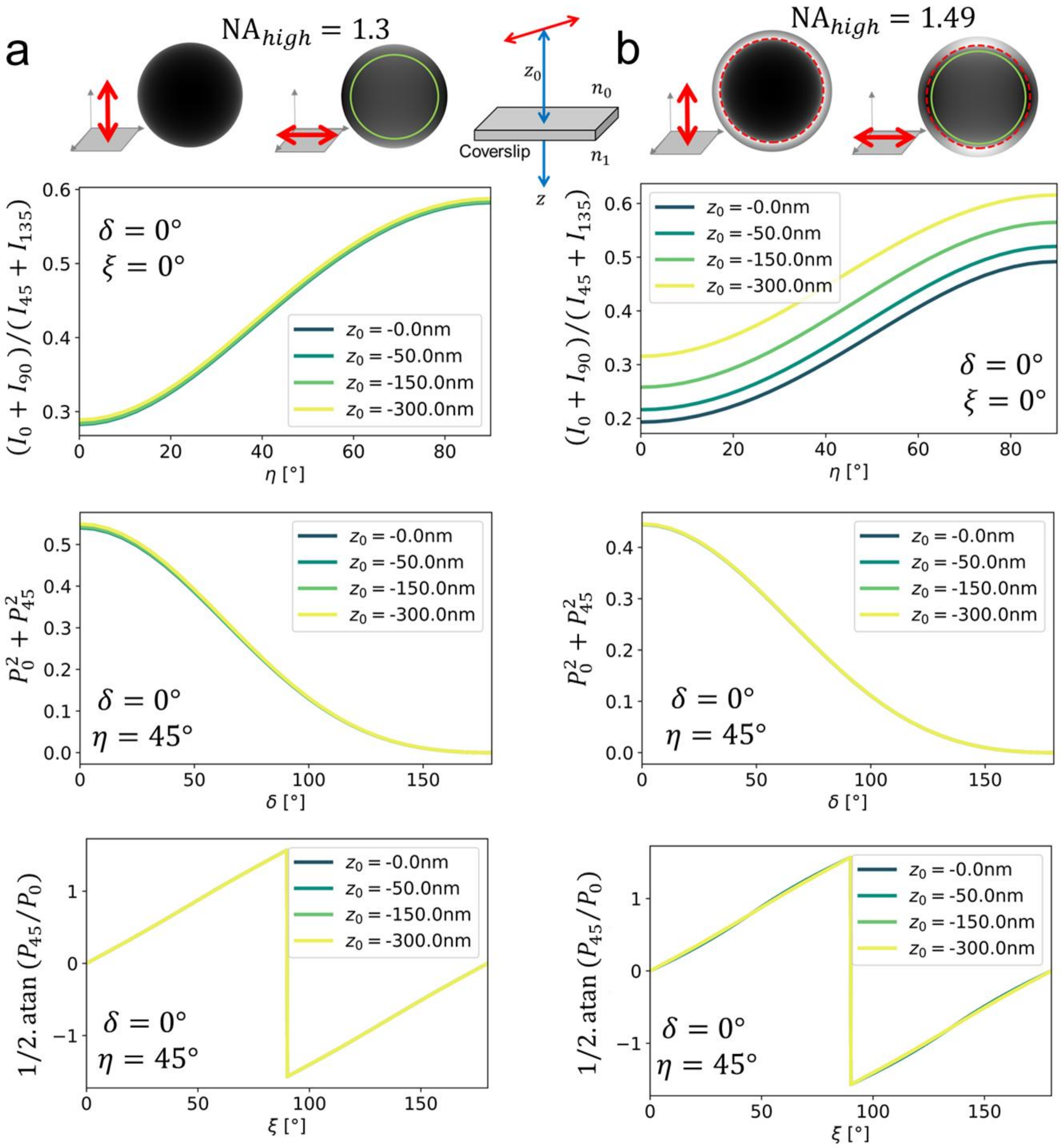


**Supplementary Fig. S2: Robustness of 4polar3D to the dipole distance to coverslip**. Conditions used for the simulations: $n_0 = 1.33$, $n_1 = 1.515$, distance to the coverslip $z_0$ variable as indicated on the graphs, defocus $z_1 = 0$. Different quantities are shown as representative of the dependence on the three parameters $(\eta, \xi, \delta)$ parameters: (upper graph) intensity ratio $(I_0 + I_{90})/(I_{45} + I_{135})$ dependence on $\eta$ ; (middle) dependence of $1/2.\,\mathrm{atan}\, P_{45}/P_0$ on ξ ; (lower graph) dependence of $P_0^2 + P_{45}^2$ on δ . For each graph, the two non-varying parameters are fixed as indicated (little dependence is seen on these parameters). Two numerical aperture conditions are displayed, keeping the ratio $NA_{low}/NA_{high} = 0.8$ constant : (a) under critical angle condition $NA_{low} = 1.04$, $NA_{high} = 1.3$; (b) critical angle condition $NA_{low} = 1.19$, $NA_{high} = 1.49$. The measured quantity $(I_0 + I_{90})/(I_{45} + I_{135})$ is seen to strongly depend on $z_0$ when $NA_{high} = 1.49$, which is due to the presence of super critical emission in the measured signal. This makes the measurement of $\eta$ highly sensitive to the dipole axial position. Setting $NA_{high}$ to an under-critical value permits to supress this dependence.

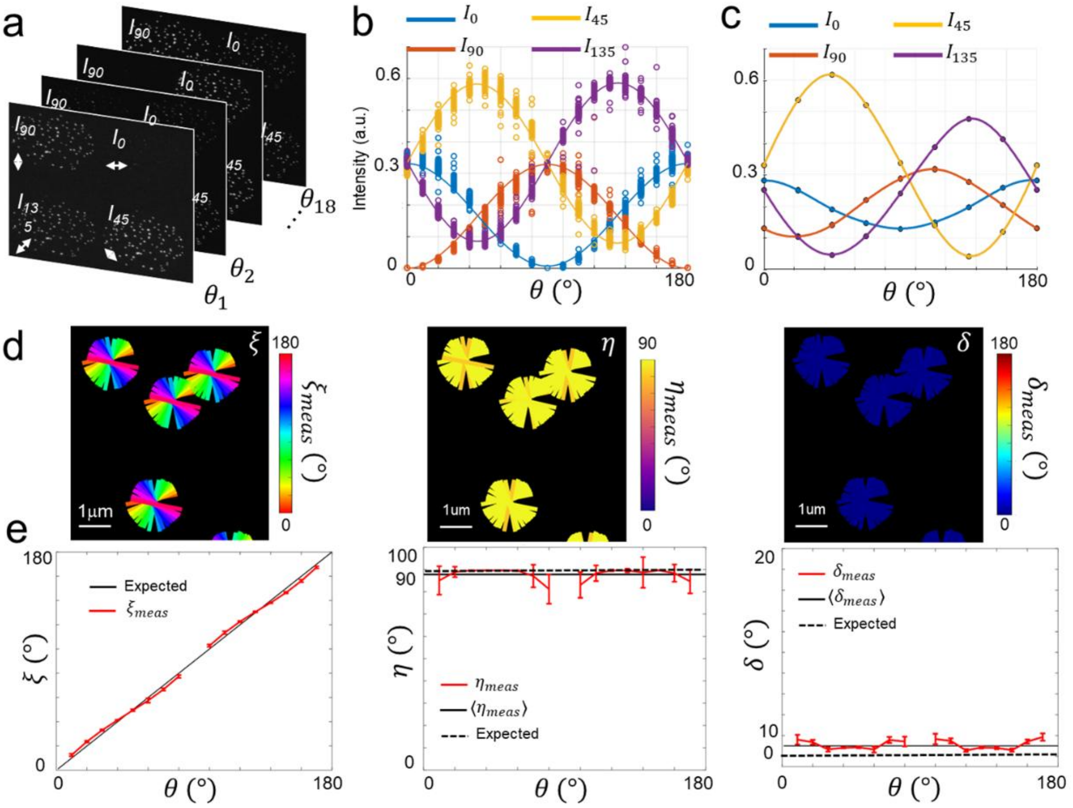


**Supplementary Figure S3. Calibration of the 4polar3D method**. (a) Recorded stack from fluorescent nanobeads followed by a rotating polarizer positioned at the pupil plane of the objective (typically, the detection polarization angle $\theta$ is rotated every 10° between 0° and 170°). (b) Recorded fluorescence from single nanobeads at the four 4polar channels, under a rotating polarizer with angle $\theta$ relative to the horizontal sample axis. Each marker represents a single nanobead, the continuous line represents the sinusoidal response fit from the averaged data. These data are used to extract the components of the $\langle K \rangle$ matrix (see Supplementary Note 2). (c) Recorded fluorescence from single nanobeads at the four 4polar channels under a rotating polarizer, when the compensating dichroic is not in place. The imbalance and low contrast of the polarized responses is the signature of diattenuation and birefringence introduced by the reflection dichroic. (d) 4polar3D stick images extracted from a stack of single nanobeads under a rotating polarizer : the orientation of the sticks is given by the retrieved angle $\xi$, their color is given respectively by $\xi$ (left), $\eta$ (middle) and $\delta$ (right). (e) Retrieved values for $\xi$ (left), $\eta$ (middle) and $\delta$ (right) as a function of the detected polarization $\theta$. The retrieved values follow the expected ones (black lines). The missing values at $\theta = 0°, 90°$ are due to the absence of detections at the (0°,90°) channels due to the true extinction obtained in these conditions ($\delta$ being close to 0°).

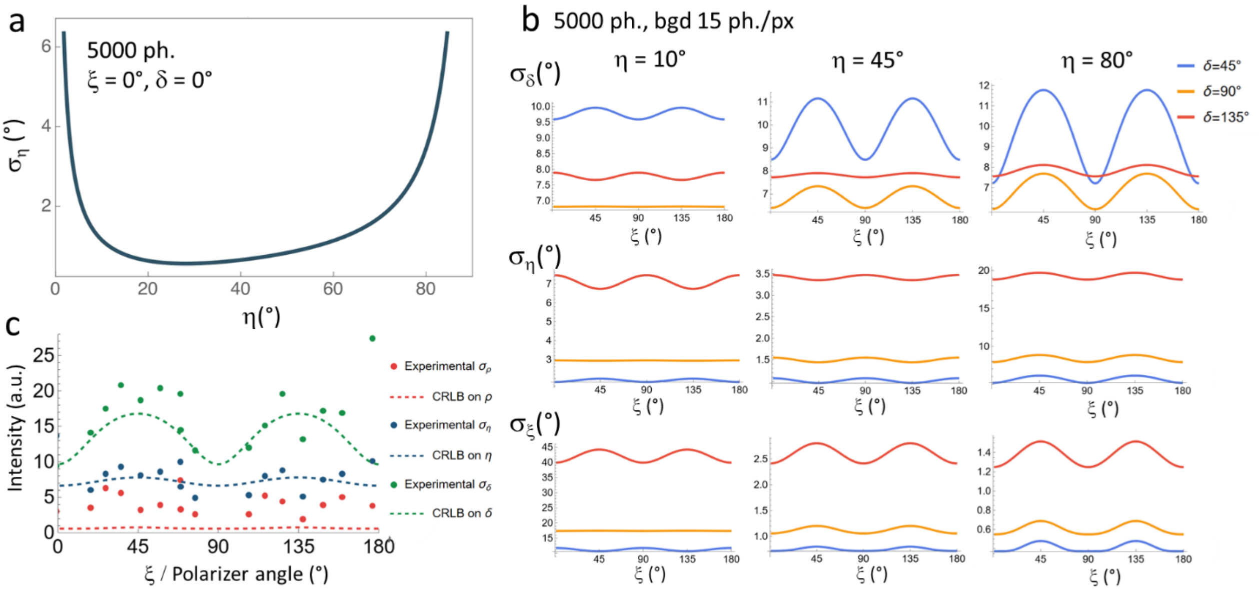


**Supplementary Fig. S4: Cramér Rao Lower Bounds (CRLB) of the 4polar3D scheme**. (a) Cramér Rao lower bond on the expectation of the $\eta$ parameter, for $\xi = 0°$ and a fixed dipole ($\delta = 0°$). Conditions used for simulations: total intensity is 5 000 photons, $n_0 = 1.33$, $n_1 = 1.515$, $z_0 = z_1 = 0$, $NA_{low} = 1.04$, $NA_{high} = 1.3$ (under critical angle condition, $NA$ ratio 0.8). (b) CRLB on the parameters $(\eta, \xi, \delta)$ calculated as a functions of ξ, for different values of $\delta$. (c) Comparison of the CRLB obtained above (dashed line) with polarized data obtained from fluorescent nanobeads (markers) (see Materials and Methods) followed by a rotating polarizer. The experimental deviations $\sigma_\eta, \sigma_\xi, \sigma_\delta$ are deduced from a measurement of 20 nanobeads.

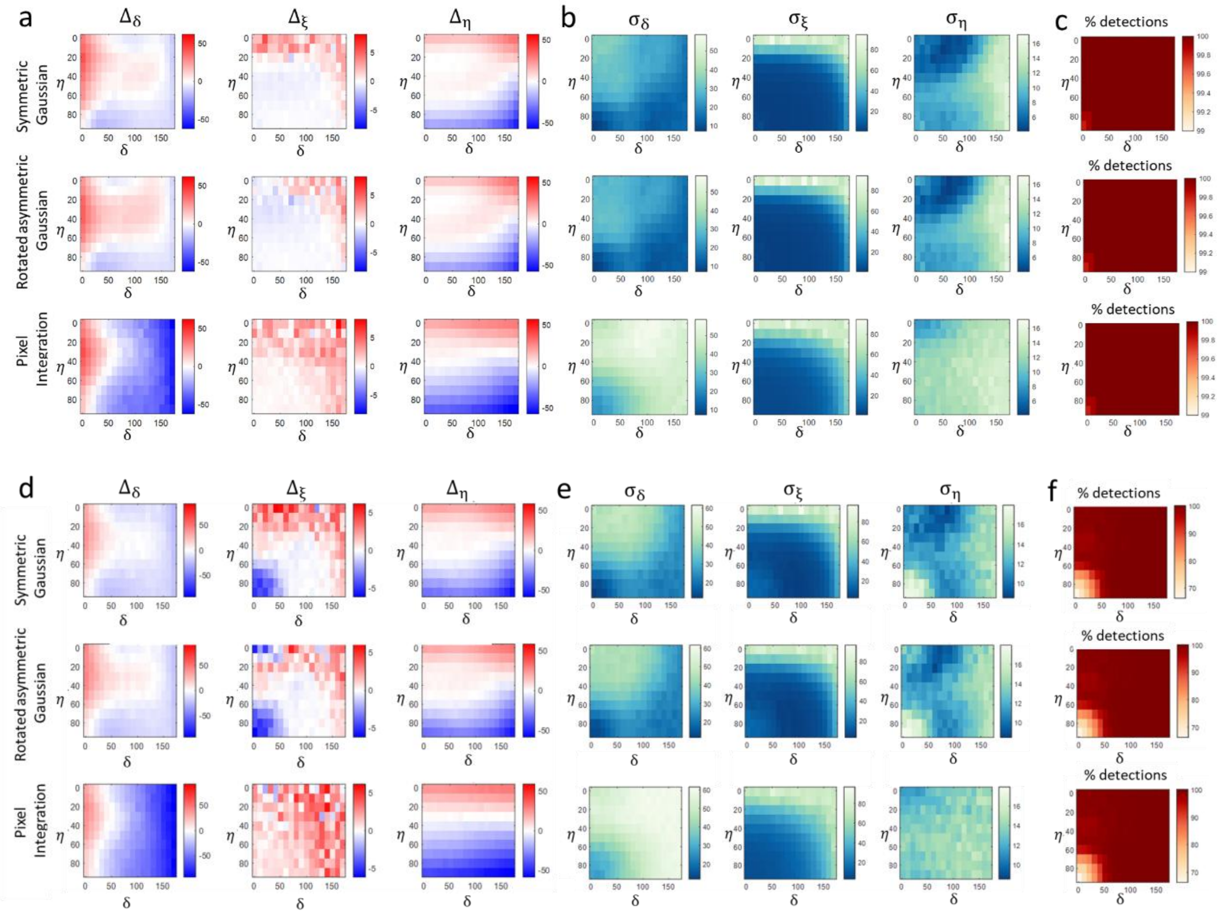


**Supplementary Fig. S5: 4polar3D Monte Carlo simulations.** Accuracy and precision (in degrees) obtained from Monte Carlo simulations (see Supplementary Note 5) on set of angles $(\eta, \xi, \delta)$ regularly spaced within the ranges $\eta = [0-90°]$, $\delta = [0-180°]$ (every 10°). $\xi$ is fixed at $\xi = 30°$ (note that the results do not strongly depend on $\xi$). Each angle set $(\eta, \xi, \delta)$ is detected for 1 000 realizations. Conditions used for simulations: Total intensity 5 000 photons (a-c) and 2 500 photons (d-f), background 10 photons/pixel, $n_0 = 1.33, n_1 = 1.515, z_0 = z_1 = 0, NA_{low} = 1.04, NA_{high} = 1.3$. For each total intensity studied, three fitting methods are displayed: symmetric Gaussian (top), rotated asymmetric Gaussian (middle), pixel integration (bottom). (a,d) Accuracy on all parameters, estimated by the mean of the differences $(\Delta_\eta, \Delta_\xi, \Delta_\delta)$ between the retrieved and the ground truth values. (b,e) precision estimated for all three parameters, using the standard deviations $(\sigma_\eta, \sigma_\xi, \sigma_\delta)$ obtained over 1000 retrievals for each condition. (c,f) Detection efficiency estimated from the % of molecules obtained from the ratio between the number of detected molecules over the initial number of molecules (1 000).

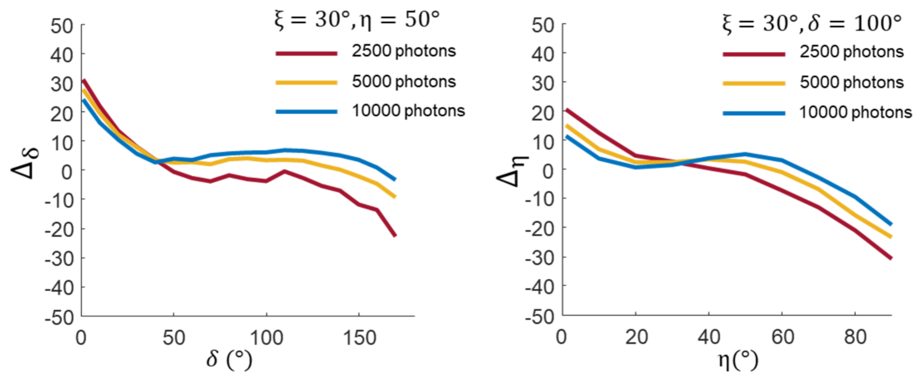


**Supplementary Fig. S6: Accuracy dependence on the molecule's total intensity,** obtained from Monte Carlo simulations. Bias $(\Delta_\eta, \Delta_\delta)$ obtained between the retrieved and the ground truth values plotted for different conditions signal (2 500, 5 000, 10 000 photons), with a background level of 10 photons/pixel, and at different (η,δ) values, for $\xi = 30°$.

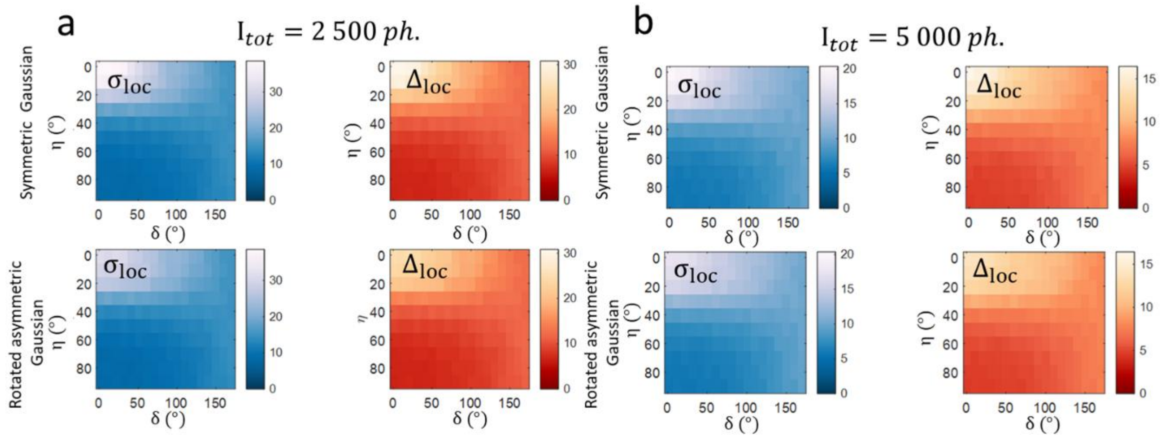


**Supplementary Fig. S7: Localization accuracy and precision**, obtained from Monte Carlo simulations. Localization accuracy $\Delta_{\mathrm{loc}}$ and localization precision $\sigma_{\mathrm{loc}}$ obtained for a set of angles regularly spaced within the ranges $\eta = [0 - 90°]$, $\delta = [0 - 180°]$, with $\xi = 30°$. Top graphs : intensities obtained by a symmetric Gaussian fit (top), lower graphs : by a rotated asymmetric Gaussian. The bias $\Delta_{\mathrm{loc}}$ is obtained by the averaged absolute value of the difference between the measured and ground truth position; the precision $\sigma_{\mathrm{loc}}$ is obtained by the standard deviation obtained on the 1 000 realizations of each angle condition (chosen to be the best precision obtained over 4 channels). Note that in these calculations, the data obtained for both $x$ and y directions are averaged. (a) condition total intensity 2 500 photons, background 10 photons/pixel, (b) total intensity 5 000 photons, background 10 photons/pixel.

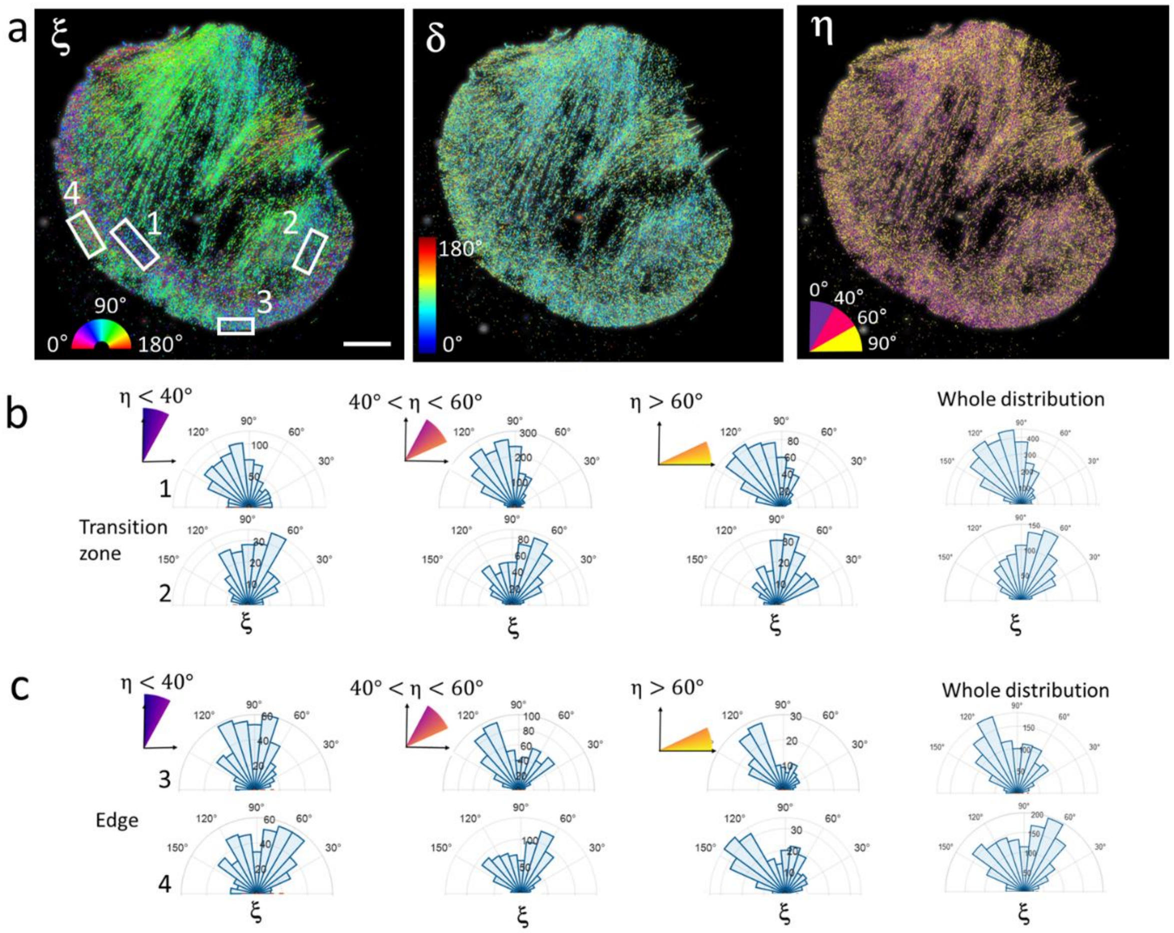


**Supplementary Fig. S8: 4polar3D in the cell lamellipodia.** (a) 4polar3D images on a fixed B16 cell labelled with phalloidin-AF568, displaying ξ (left), $\delta$ (middle) and η (right). η is shown using a binned colormap with indicated angular sectors. (b) Polar histograms of ξ for given populations of η values : off-plane ($\eta < 40°$), intermediate ($40° < \eta < 60°$), in-plane ($\eta > 60°$), as well as the whole distribution, in different ROIs (of typically 1x1 $\mu m^2$) in the transition zone of the cell. (c). Similar distributions, at the edge of the cell within the lamellipodium. While the transition zones display quite continuous, broad but directional distribution not dependent on the off-plane tilt angle of the molecules, the edge distributions are quite variable and often display bimodal distributions with more or less pronounced lobes. The bimodal features are more visible for in-plane filaments. Scale bar : 5$\mu$m (a).

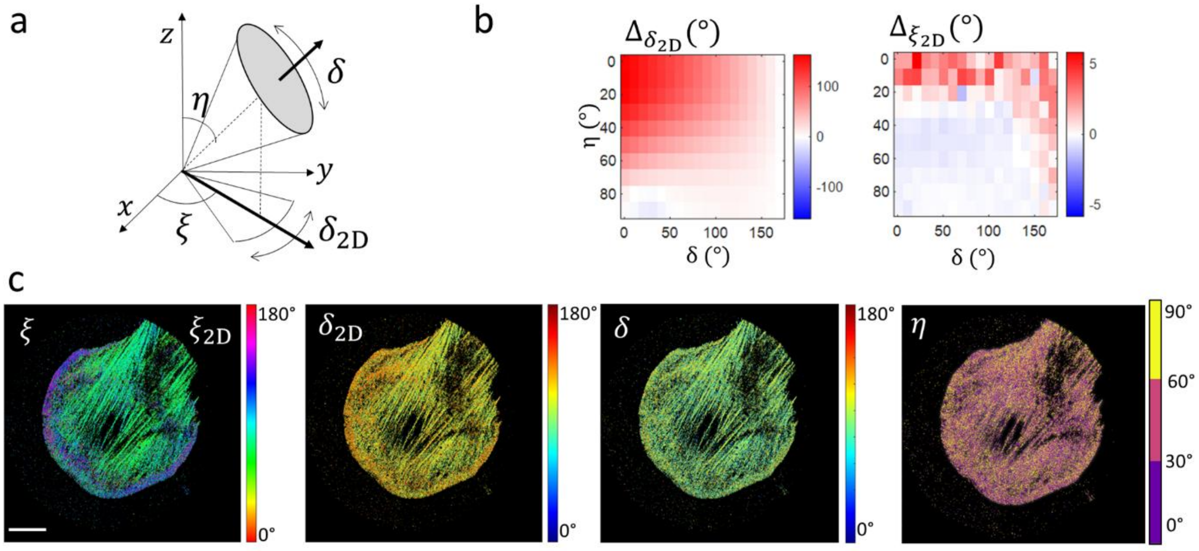

**Supplementary Fig. S9: Estimation bias of 4polar2D versus 4polar3D.** (a) Schematic representation of the 2D projected wobbling quantity $\delta_{2D}$, estimated by the method 4polar2D[7]. (b) Theoretical bias induced by a 2D measurement in 4polar2D (i.e. ignoring the estimation of the $z$ dipole contributions (Supplementary Note 1). The estimated wobbling angle $\delta_{2D}$ and in-plane orientation $\xi_{2D}$ are estimated using the 4polar (2D) formalism as previously developed[7] and compared to the ground truth using bias quantities : $\Delta_{\delta_{2D}} = \delta_{2D} - \delta$ and $\Delta_{\xi_{2D}} = \xi_{2D} - \xi$. The bias is averaged over 1 000 realizations, from a Monte Carlo simulation of single molecule images spanning all possible $(\eta, \delta)$ values every 10°, with $\xi = 30°$. A total intensity of 5 000 photons and a background of 10 photons/pixel are used. (c) 4polar (2D) and 4polar3D approaches applied to the analysis of 4polar3D data performed on a B16 cell displaying a wide lamellipodium. Left : $\xi$ (both $\xi_{2D}$ and $\xi$ lead to similar images). Middle : $\delta_{2D}$ and $\delta$. The estimation of $\delta_{2D}$ clearly leads to an overestimation of the wobbling in the lamellipodium, as compared with the 3D analysed $\delta$ values. Right : $\eta$. High $\delta_{2D}$ values are visualized in region of off-planes molecules ($\eta < 40°$). Scale bar : 5 µm (c).

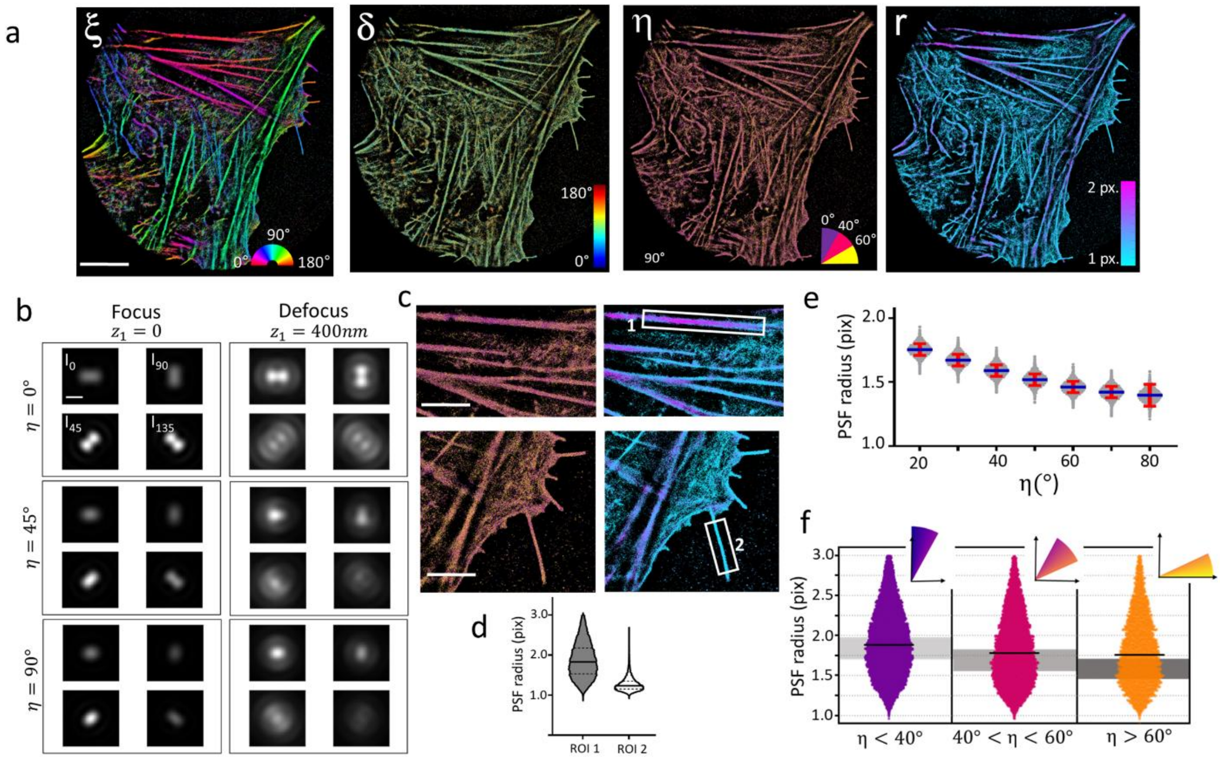


**Figure S10. 4polar3D on actin stress fibers.** (a) 4polar3D images on a fixed U2OS cell labelled with phalloidin-AF568, displaying from left to right: ξ, $\delta$, η and r (PSF radius, in pixels). η is shown using a binned colormap with indicated angular sectors. (b) Modelled 4polar3D PSFs for different situations of η angles and defocus $z_1$ ($\xi = 30°$, $\delta = 100°$) (parameters : $n_0 = 1.33$, $n_1 = 1.515$, $NA_{low} = 1.1$, $NA_{high} = 1.3$, distance to the coverslip $z_0 = 0$). (c) zoomed η and r images of different regions of (a). (d) Values of r represented for two regions of interest (ROI) of (c) displayed as rectangles in (c) (30 000 molecules (ROI1) and 100 000 molecules (ROI2) are detected). (e) PSF radius r values retrieved from theoretical PSFs in noise conditions (5 000 photons, 10 background photon/pixel) for PSF generated from random ξ, $\delta = 100°$, and for different values of η. (1 000 PSFs are analyzed per η value). Averaged values or r are shown in red. (f) PSF radius r obtained from different sub-populations of η (as indicated) retrived from molecules detected in ROI1. Grey regions of r indicate the range of r expected for in-focus molecules in the corresponding range of η. Scale bars : 10 µm (a), 800 nm (b), 3 µm (c).
